# Slow oscillation-spindle coupling predicts sequence-based language learning

**DOI:** 10.1101/2020.02.13.948539

**Authors:** Zachariah R. Cross, Randolph F. Helfrich, Andrew W. Corcoran, Adam J. O. Dede, Mark J. Kohler, Scott W. Coussens, Lena Zou-Williams, Matthias Schlesewsky, M. Gareth Gaskell, Robert T. Knight, Ina Bornkessel-Schlesewsky

## Abstract

Sentence comprehension involves the rapid decoding of both semantic and grammatical information, a process fundamental to communication. As with other complex cognitive processes, language comprehension relies, in part, on long-term memory. However, the electrophysiological mechanisms underpinning the initial encoding and generalisation of higher-order linguistic knowledge remain elusive, particularly from a sleep-based consolidation perspective. One candidate mechanism that may support the consolidation of higher-order language is the temporal coordination of slow oscillations (SO) and sleep spindles during non-rapid eye movement sleep (NREM). To examine this hypothesis, we analysed electroencephalographic (EEG) data recorded from 35 participants (M_age_ = 25.4, SD = 7.10; 16 males) during an artificial language learning task, contrasting performance between individuals who were given an 8hr nocturnal sleep period or an equivalent period of wake. We found that sleep relative to wake was associated with superior performance for rules that followed a sequence-based word order. Post-sleep sequence-based word order processing was further associated with less task-related theta desynchronisation, an electrophysiological signature of successful memory consolidation, as well as cognitive control and working memory. Frontal NREM SO-spindle coupling was also positively associated with behavioural sensitivity to sequence-based word order rules, as well as with task-related theta power. As such, theta activity during retrieval of previously learned information correlates with SO-spindle coupling, thus linking neural activity in the sleeping and waking brain. Taken together, this study presents converging behavioral and neurophysiological evidence for a role of NREM SO-spindle coupling and task-related theta activity as signatures of successful memory consolidation and retrieval in the context of higher-order language learning.

**SIGNIFICANCE STATEMENT:** The endogenous temporal coordination of neural oscillations supports information processing during both wake and sleep states. Here we demonstrate that slow oscillation-spindle coupling during non-rapid eye movement sleep predicts the consolidation of complex grammatical rules and modulates task-related oscillatory dynamics previously implicated in sentence processing. We show that increases in theta power predict enhanced sensitivity to grammatical violations after a period of sleep and strong slow oscillation-spindle coupling modulates subsequent task-related theta activity to influence behaviour. Our findings reveal a complex interaction between both wake- and sleep-related oscillatory dynamics during the early stages of language learning beyond the single word level.

## Introduction

The human brain is adept at extracting regularities from sensory input, a process pivotal for generating knowledge of one’s physical and social environment (Santolin & Saffran, 2018). Notably, learning of such regularities plays a key role in the development of linguistic competencies, enabling the implicit acquisition of grammatical rules embedded in ambient speech (Cross et al., 2021; Isbilen et al., 2022; Romberg & Saffran, 2010, 2010). While this perspective of language learning has informed insights concerning the encoding of local dependencies, the acquisition of more complex linguistic structures remains less understood. Here, we address this gap from the perspective of sleep-based memory consolidation, a well-established mechanism governing the generalisation of knowledge from sensory experience (Brodt et al., 2023; Diekelmann et al., 2009; Xie et al., 2018).

A plethora of evidence (for review, see Rasch and Born 2013) demonstrates that sleep plays an active role in memory by consolidating and generalising mnemonic information. This dynamic account of the sleeping brain is captured by the Active System Consolidation hypothesis (ASC; (Born & Wilhelm, 2012; Klinzing et al., 2019). Core to ASC is that sleep facilitates repeated reactivation of encoded memory representations (Rasch & Born, 2013). This reactivation is dependent on cortical glutamatergic synapses, which weaken during prolonged wakefulness (Kavanau, 1997; Rasch & Born, 2013). The ASC is supported by electrophysiological evidence that learned sequences are replayed during non-rapid eye-movement (NREM) sleep, potentially via sleep spindle and slow oscillatory (SO) activity. Sleep spindles are bursts of electrical activity occurring between 11 – 16 Hz, while SOs centred at 1 Hz reflect synchronized membrane potential fluctuations between hyperpolarised up-states and depolarised down-states of neocortical neurons (Crunelli & Hughes, 2010; Vyazovskiy & Harris, 2013). The precise coupling between SOs and spindles provides a temporal receptive window for the replay of hippocampal memory traces and their transfer to cortex for long-term storage (Bastian et al., 2022; Mikutta et al., 2019). Critically, the transfer of newly encoded information from hippocampus to cortex enables generalisation of mnemonic information, allowing cortex to learn the regularities of sensory input gradually – a process known to support language learning (Cross et al., 2018; Davis & Gaskell, 2009; Rasch, 2017).

Mechanisms of sleep-based memory consolidation have been associated with aspects of language learning, including novel-word learning (Bakker et al., 2015; James et al., 2017; Mirković & Gaskell, 2016) as well as the generalisation of grammatical rules (Batterink et al., 2014; Nieuwenhuis et al., 2013). Positive associations have also been identified between rapid eye-moment (REM) sleep percentages and language learning proficiency (De Koninck et al., 1989, 1990), supporting a link between REM sleep and language learning. To elucidate the mechanism of this relationship, Thompson et al. (2021) examined oscillatory dynamics during REM sleep and demonstrated that sleep spindles and theta power predicted language learning among individuals engaged in second-language immersion programs. This effect was stronger when time-locked to eye movements during REM sleep.

Together, extant work on sleep and language learning underscore the significance of both REM and NREM sleep, sleep spindles, and theta power in facilitating second language learning. However, work examining the association between sleep and language often involves only behavioural measures as proxies for memory consolidation (e.g., Mirković & Gaskell, 2016; Nieuwenhuis et al., 2013), or examines structure (e.g., grammar; Nieuwenhuis et al., 2013) and meaning (i.e., semantics; Bakker et al., 2015; Batterink et al., 2017; Batterink & Paller, 2017) in the language input separately (cf. Batterink et al., 2014). Markers of sleep-based memory consolidation are also often based on coarse experimental contrasts (i.e., sleep vs. wake conditions) or macroarchitectural measures (i.e., percent time spent in a particular sleep stage), rather than neurophysiological events that can more directly test models of systems consolidation anchored in NREM sleep, such as SO-spindle coupling. Online EEG measures during language learning and comprehension and their relation to offline states, such as sleep, are also lacking.

From this perspective, neurobiological models of sleep, memory, and language processing would benefit from a direct investigation of the relation between sleep and higher-order language, such as at the sentence level that have differing grammatical rules (Cross et al., 2018; Rasch, 2017; Schreiner & Rasch, 2017), in conjunction with online measures of neural activity. This would extend our understanding of the complexity of language learning beyond single words, and how the generalisation of newly acquired linguistic knowledge is supported by sleep (for review, see Cross et al., 2018) and how the brain learns environmental regularities that span multiple scales of complexity and how this information is organised across sleep and wake.

Here, we present data addressing the contribution of sleep-based memory consolidation to complex rule learning in language at the sentence level. We used the modified miniature language Mini Pinyin (Cross et al., 2021), which is modelled on Mandarin Chinese, to contrast rules that instantiate a fixed or flexible word order. Mandarin naïve Monolingual native English speakers completed a learning task where they were shown pictures of two-person events, followed by a sentence describing the event in the picture. During this task, participants learned varying word order rules without explicit instruction and then completed a baseline memory task prior to either 8hr of sleep or an equivalent period of wake (Figure 1). Participants then completed a delayed memory task to assess changes in memory of the word order rules after the 8hr delay.

**Figure 1.**
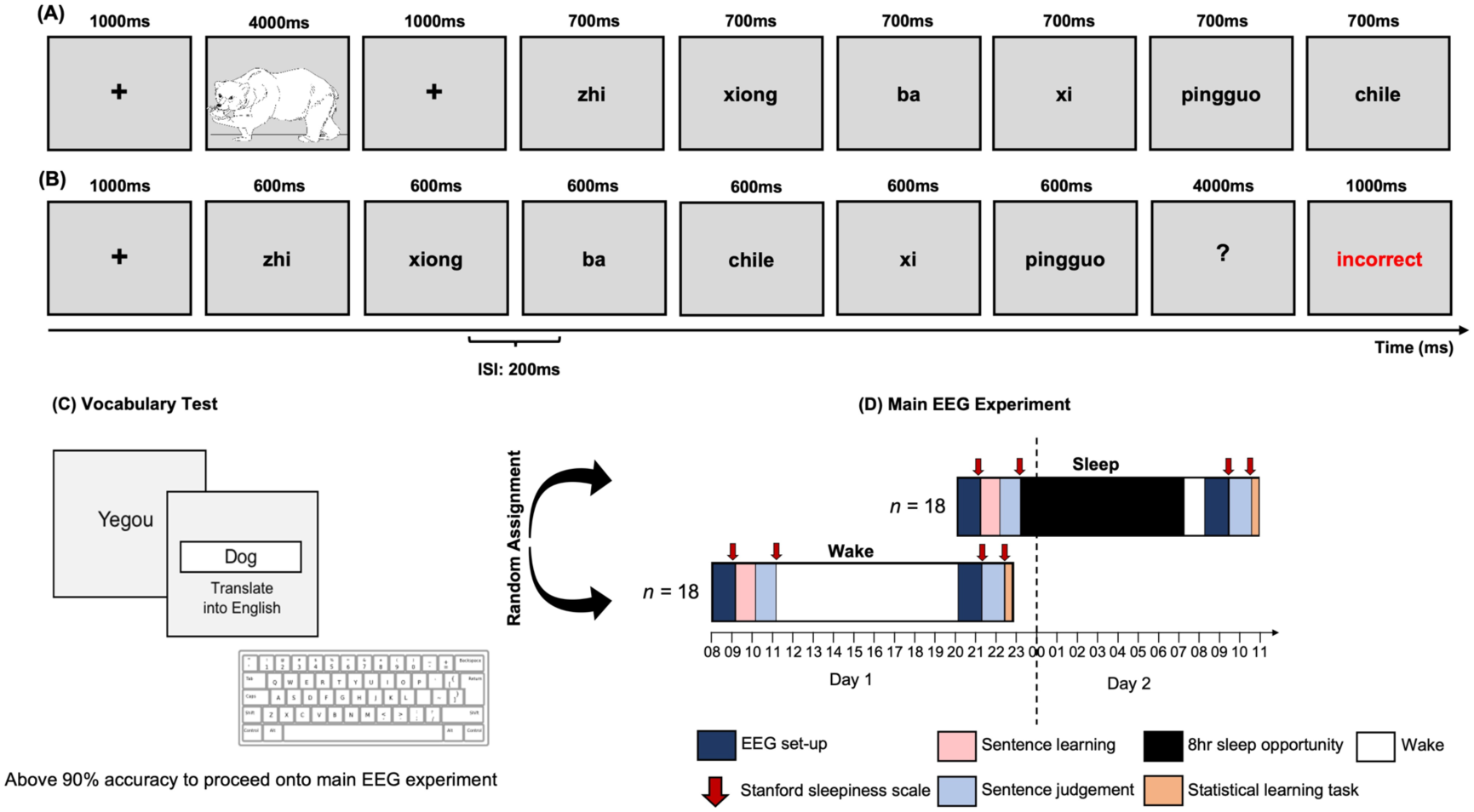
Illustration of stimulus presentation and experimental protocol. **(A)** Schematic representation of a single trial of a grammatical sentence during the sentence learning task. **(B)** Schematic representation of a single trial during the baseline sentence judgement task. This sentence is a violation of the verb-position, whereby the verb *chile* is positioned in the middle of the sentence when it should be positioned at the end of the sentence. Here, the participant incorrectly categorised this sentence as grammatical, and thus received feedback indicating that their response was incorrect. **(C)** Schematic diagram of the vocabulary test, which required participants to translate the nouns (e.g., *yegou*) into English (e.g., *dog*) using a keyboard. **(D)** Experimental protocol representing the time course of the conditions (sleep, wake) and testing sessions (sentence learning task, baseline, and delayed sentence judgement tasks). After completing the vocabulary test, participants were randomly assigned to either the sleep or wake conditions, with each participant only completing one of the two conditions. Time is represented along the x-axis, while each coloured block corresponds to a different task during the experimental protocol.

We focussed on theta oscillations (∼ 3 – 7 Hz), which were quantified using complex Morlet wavelets across sentence presentation during the memory tasks. Theta oscillations are implicated in relational binding and memory-based decision making (Backus et al., 2016; Buzsáki, 2002; Jacobs et al., 2006). From this perspective, theta should track successful language learning and sleep-based consolidation (Cross et al., 2018). We further quantified whole-scalp NREM SO-spindle coupling by detecting spindle events and quantifying the percentage of spindle events that occurred during SO events. SO-spindle coupling as well as task-related theta power were used to independently predict language learning, and to determine whether task-related theta is modulated by sleep-based memory consolidation.

## Methods

### Participants

We recruited 36 right-handed participants who were healthy, monolingual, native English-speakers (16 male) aged 18 – 40 years old (M_age_ = 25.4, SD = 7.0). Participants were randomly assigned to either a Sleep (*n* = 18) or Wake condition. All participants reported normal or corrected-to-normal vision, no history of psychiatric disorders, substance dependence, or intellectual impairment, and were not taking medication that influenced sleep or neuropsychological measures. All participants provided informed consent and received a $120 honorarium. One participant from the Sleep condition was removed from the analysis due to technical issues during the experimental tasks and sleep period, resulting in a total sample size of 35 (M_age_ = 25.4, SD = 7.10; 16 males; Sleep *n* = 17). Ethics approval was granted by the University of South Australia’s Human Research Ethics committee (I.D: 0000032556).

### Screening and control measures

The Flinders Handedness Survey (FLANDERS; Nicholls et al., 2013) was used to screen handedness, while the Pittsburgh Sleep Quality Index (PSQI; Buysse et al., 1989) screened for sleep quality. PSQI scores ranged from 1-5 (*M* = 2.9, *SD* = 1.33) out of a possible range of 0 – 21, with higher scores indicating worse sleep quality. Prospective participants with scores > 5 were unable to participate. As an additional control, the Stanford Sleepiness Scale (SSS) was administered at the beginning and end of the experiment to measure self-perceived sleepiness.

### Electroencephalography

The electroencephalogram (EEG) was recorded during the learning and sentence judgement tasks and sleep opportunities using a 32-channel BrainCap with sintered Ag/AgCI electrodes (Brain Products, GmbH, Gilching, Germany) mounted according to the extended International 10-20 system. The reference was located at FCz, with EEG signals re-referenced to linked mastoids offline. The ground electrode was located at AFz. The electrooculogram (EOG) was recorded via electrodes located 1cm from the outer canthus of each eye (horizontal EOG) and above and below participants’ left eye (vertical EOG). Sub-mental electromyography (EMG) was added to facilitate accurate scoring of sleep periods. The EEG was amplified using a BrainAmp DC amplifier (Brain Products GmbH, Gilching, Germany) using an initial band-pass filter of DC – 250 Hz with a sampling rate of 1000 Hz.

### Vocabulary and structure of Mini Pinyin

Stimuli consisted of sentences from a modified miniature language based on Mandarin Chinese (Cross et al., 2021). This language contained 32 transitive verbs, 25 nouns, 2 coverbs, and 4 classifiers. The nouns included 10 human entities, 10 animals and 5 objects (e.g., *apple*). Each category of noun was associated with a specific classifier, which always preceded each of the two noun phrases in a sentence. As illustrated in Figure 2B, *ge* specifies a human noun, *zhi* for animals, and *xi* and *da* for small and large objects, respectively. Overall, this stimulus set contained 576 unique sentences (288 grammatical, 288 ungrammatical) which are divided into two equivalent sets (see Cross et al., 2021) for a complete description of the stimuli; for the complete set of stimuli, visit: https://tinyurl.com/3an438h2).

**Figure 2.**
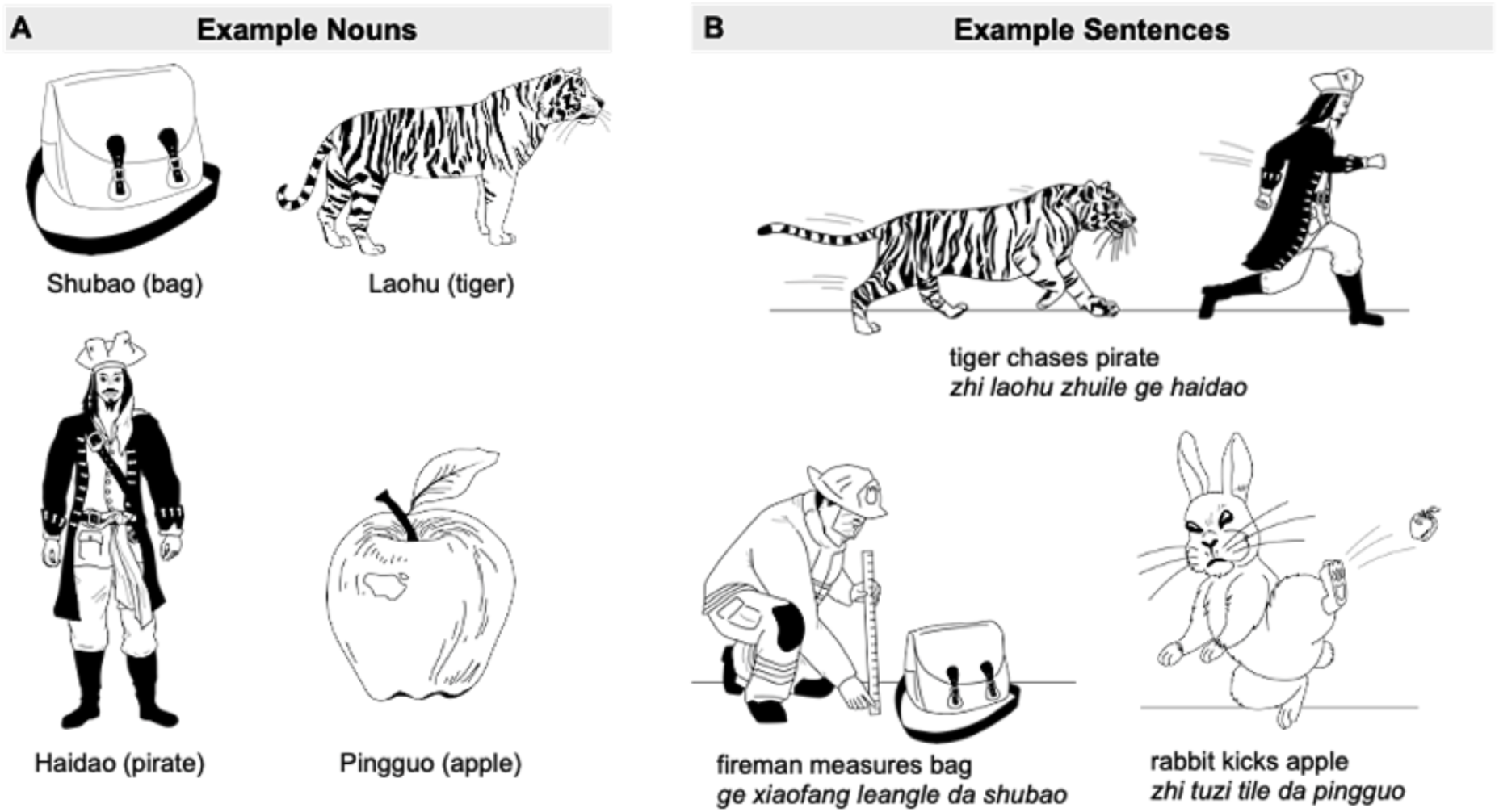
Example of images used in vocabulary and sentence learning phases. **(A)** Portion of the 25 illustrations used in the vocabulary booklet, which included human, animal, and inanimate objects (i.e., bag, apple). **(B)** Portion of the illustrations used in the sentence learning task, illustrating the interaction between two entities. Note that the entities used in sentence learning task are the same as the illustrations used in the vocabulary booklet.

We focussed on a subset of sentence conditions to investigate the mechanisms underlying the learning of different word order rules, which fundamentally differs between natural languages (for review, see Bates et al., 2001). Languages like English and Dutch rely primarily on word order, while languages like German and Turkish rely more on cues such as case marking and animacy (Bornkessel & Schlesewsky, 2006; Bornkessel-Schlesewsky et al., 2015; MacWhinney et al., 1984). From this perspective, Mini Pinyin enabled a comparison between sentences with differing word orders (see Figure 3A), and the influence sleep may have on the respective consolidation of fixed and flexible word order rules. The subset of stimuli in the current analysis contained 96 sentences in the sentence learning task and 144 sentences in the grammaticality judgement tasks. The remaining sentences were considered fillers. These filler sentences included sentences that violated classifier-noun pairs, and thus were not suitable for testing predictions regarding fixed and flexible word order processing (for a full description of all sentence conditions present in this language, please see (Cross et al., 2021).

**Figure 3.**
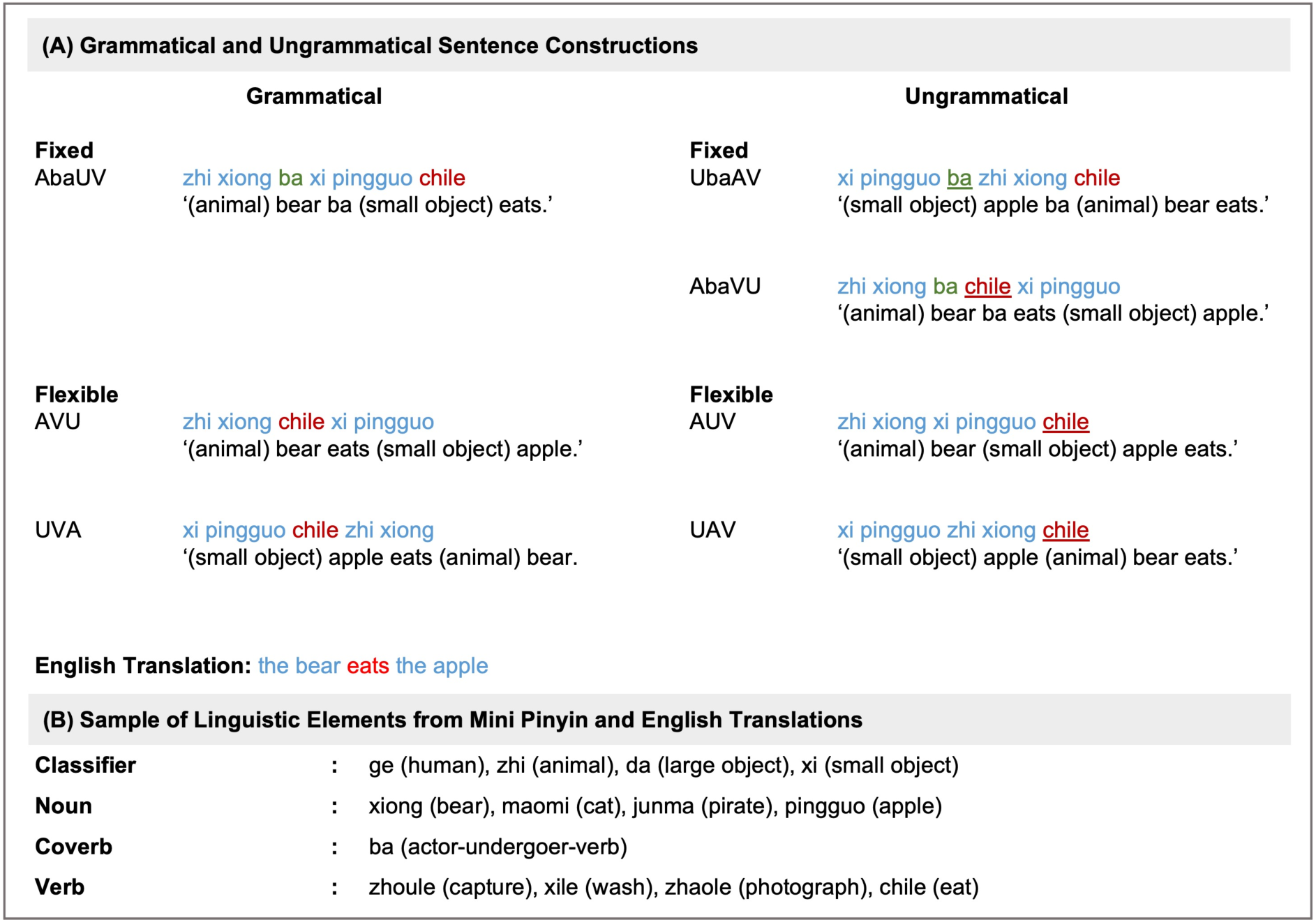
Exemplar word order rules and vocabulary items of Mini Pinyin. **A)** Example of grammatical and ungrammatical fixed and flexible word order sentences. Classifiers and nouns are coded in blue, while verbs are red. The coverb *ba* is coded in green. For the ungrammatical sentences (right), the point of violation in the sentence is underlined. The direct English translation for each sentence construction is provided below (i.e., *the bear eats the apple*). **(B)** A sample of the linguistic elements present in Mini Pinyin and their English translation. Note that *ba* does not have a specific meaning, but when present in a sentence, instantiates a strict actor-undergoer-verb word order.

As is apparent in Figure 3A, sentences that do not contain the coverb *ba* (i.e., actor-verb-undergoer, AVU; undergoer-verb-actor, UVA) yield a flexible word order, such that understanding *who is doing what to whom* is not dependent on the ordering of the noun phrases. Instead, determining *who is doing what to whom* is facilitated by animacy cues. For instance, in the UVA condition, *the bear* is interpreted as the actor despite the first noun phrase being *the apple*, since it is implausible for an apple to eat a bear. Therefore, both AVU and UVA are grammatical constructions. By contrast, sentences such as A*ba*UV yield a fixed word order, such that the inclusion of *ba* strictly renders the first noun phrase as the actor. Note that the positioning of the verb is critical in sentences with and without a coverb. With the inclusion of a coverb, the verb must be placed at the end of the sentence, while the verb must be positioned between the noun phrases in constructions without a coverb.

### Experimental protocol

Participants received a paired picture-word vocabulary booklet containing the 25 nouns and were asked to maintain a minimum of 7hrs sleep per night (see Figure 2A for a portion of nouns from the vocabulary booklet). Participants were required to learn the 25 nouns to ensure that they had a basic vocabulary of the nouns to successfully learn the 32 transitive verbs. They were asked to record periods of vocabulary learning in an activity log. Participants were instructed to study the booklet for at least fifteen minutes per day and were informed that they would need to pass a vocabulary test before commencing the main experimental protocol. After approximately one week, participants returned to complete the main experimental session, where EEG was recorded during a sentence learning task, baseline, and delayed sentence judgement tasks.

### Vocabulary test

Participants completed a vocabulary test by translating the nouns from Mini Pinyin into English using a keyboard, as illustrated in Figure 1C. Each trial began with a 600ms fixation cross, followed by the visual presentation of the noun for up to 20s. Prospective participants who scored < 90% were unable to complete the main experimental EEG session. As such, all 36 participants included in the current paper obtained over 90% correct on the vocabulary test. The proportion of individuals who did not pass the vocabulary test was small (e.g., approximately less than 5 cases); however, the exact number was not recorded.

### Sentence learning

Sentence and picture stimuli were presented using OpenSesame (Mathôt et al., 2012). During sentence learning, pictures were used to depict events occurring between two entities. The pictures and entities shown during the learning task were combinations of the static pictures shown in the vocabulary booklet (for an example of booklet versus sentence learning picture stimuli, see Figure 2A and 2B, respectively).

While participants were aware that they would complete sentence judgement tasks at a later point, no explicit description of or feedback regarding grammatical rules was provided during the learning task. Each picture corresponded to multiple sentence variations, similar to the grammatical conditions in Figure 3A. Picture-sentence pairs were presented to participants as correct language input. Participants were presented with a fixation cross for 1000ms, followed by the picture illustrating the event between two entities for 5000ms. A sentence describing the event in the picture was then presented on a word-by-word basis. Each word was presented for 700ms followed by a 200ms ISI. This pattern continued for the 96 reported combinations, until the end of the task, which took approximately 40 minutes. The 96 sentences included in this analysis included the flexible (i.e., AVU, UVA) and fixed (i.e., AbaUV) sentence constructions. Sentences considered as fillers contained a coverb that was not *ba*, and thus were not relevant to testing the predictions posited in the current analysis. During this task, participants were required to learn the structure of the sentences and the meaning of the verbs, classifiers and the coverb *ba*. Stimuli were pseudo-randomised, such that no stimuli of the same construction followed each other, and each sentence contained a different combination of nouns and verbs. This was done to encourage learning of the underlying grammatical rules rather than episodic events of individual sentences. Further, the two lists of sentences were counterbalanced across participants and testing session. Following the sentence learning task, participants completed the baseline judgement task.

### Baseline and delayed judgement tasks

The baseline sentence judgement task taken immediately after learning provided a baseline to control for level of encoding, while the delayed judgement task took place ∼12hrs after the learning and baseline judgement tasks. During both judgement tasks, 288 sentences without pictures (144 grammatical, 144 ungrammatical), 156 of which are reported here, were presented word-by-word with a presentation time of 600ms and an ISI of 200ms. The 156 included sentences included a combination of grammatical and ungrammatical flexible and fixed sentence constructions, while the 132 sentences that were considered fillers contained coverbs that were not *ba*, and classifier-noun pair violations, and thus were not relevant to testing the predictions of the current analysis. Participants received feedback on whether their response was correct or incorrect during the baseline but not the delayed judgement task. This was to ensure that participants were able to continue learning the language without explicit instruction. Figures 1A and 1B illustrate the sequence of events in the sentence learning and baseline judgement tasks, respectively.

Participants were instructed to read all sentences attentively and to judge their grammaticality via a button-press. As a cue for judgment, a question mark appeared in the centre of the monitor for 4000ms after the offset of the last word. Two lists of sentence stimuli were created, which were counterbalanced across participants and the baseline and delayed sentence judgement tasks. Half of the sentences were grammatical, with each of the grammatical constructions shown an equal number of times. The other half of the sentences were ungrammatical constructions. Stimuli were pseudo-randomised, such that no stimuli of the same construction followed each other.

### Main experimental procedure

For the wake condition, participants completed the vocabulary test and EEG setup at ∼08:00hr. The learning task was administered at ∼09:00hr, followed by the baseline judgement task, with EEG recorded during both the learning and judgement task. Participants then completed the behavioural control tasks and were free to leave the laboratory to go about their usual daily activities, before returning for EEG setup and the delayed judgement task at ∼21:00hr the same day. EEG was also recorded during the delayed judgement task.

Participants in the sleep condition arrived at ∼20:00hr to complete the vocabulary test and EEG setup before completing the learning task at ∼21:00hr, followed by the baseline judgement task, with EEG recorded during both the learning and judgement tasks. Participants were then given an 8hr sleep opportunity from 23:00hr – 07:00hr. Polysomnography was continuously recorded and later scored. After waking, participants were disconnected from the head box and given a ∼1hr break to alleviate sleep inertia before completing the delayed judgement task and behavioural control tasks. During this time, participants sat in a quiet room and consumed a small meal. Resting-state EEG recordings were obtained during quiet sitting with eyes open and eyes closed for two minutes, respectively. See Figure 1D for a schematic of the experimental protocol.

## Data Analysis

### Behavioural analysis

Two measures of behavioural performance were calculated. For the behavioural analysis, grammaticality ratings were calculated on a trial-by-trial basis, determined by whether participants correctly identified grammatical and ungrammatical sentences. For EEG analyses, memory performance was quantified using the sensitivity index (d’) from signal detection theory (Stanislaw & Todorov, 1999). Hit Rate (HR) and False Alarm rate (FA) were computed to derive d’, defined as the difference between the z transformed probabilities of HR and FA (i.e., d’ = z[HR] – z[FA]), with extreme values (i.e., HR and FA values of 0 and 1) adjusted using the recommendations of (Hautus, 1995).

### EEG recording and pre-processing

Task-related EEG analyses during the baseline and delayed sentence judgement tasks were performed using MNE-Python (Gramfort et al., 2013). EEG data (C3, C4, CP1, CP2, CP5, CP6, Cz, F3, F4, F7, F8, FC1, FC2, FC5, FC6, Fp1, Fp2, Fz, O1, O2, P3, P4, P7, P8, Pz) were re-referenced offline to the average of both mastoids and filtered with a digital phase-true finite impulse response (FIR) band-pass filter from 0.1 – 40 Hz to remove slow signal drifts and high frequency activity. Data segments from -0.5 – 6.5s relative to the onset of each sentence were extracted and corrected for ocular artefacts using Independent Component Analysis (fastica; (Hyvarinen, 1999). Epochs were dropped when they exceeded a 150 μV peak-to-peak amplitude criterion or were identified as containing recordings from flat channels (i.e., < 5 μV).

#### Task-related time frequency analysis

To determine the individualised ranges used to define the theta frequency band, individual alpha frequency (IAF) was estimated from participants’ pre- and post-experiment resting-state EEG recording. IAFs were estimated from an occipital-parietal cluster (P3/P4/O1/O2/P7/P8/Pz/Oz) using philistine.mne.savgol_iaf (see Corcoran et al., 2018) implemented in MNE (philistine.mne). IAF-adjusted frequency bandwidths were calculated according to the harmonic frequency architecture proposed by (Klimesch, 2012, 2013) and which is in line with previous work (Corcoran et al., 2018; Cross et al., 2022; Doppelmayr et al., 1998; Sauppe et al., 2021), in which the centre frequency of each successive band constitutes a harmonic series scaled in relation to the IAF.

We conducted task-related time-frequency analyses by convolving the pre-processed EEG with a family of complex Morlet wavelets using the MNE function tfr_morlet. Theta activity was analysed using wavelet cycles, with the mother wavelet defined as the centre frequency value divided by four. Relative power change values in the post-stimulus interval were computed as a relative change from a baseline interval spanning -0.5s to the onset of each sentence. As such, theta power during the sentence period reflects deviations from the baseline interval, such that higher theta power would indicate an increase in power relative to baseline, while a decrease in power indicates a decrease in power relative to baseline. 500ms was added to the beginning and end of each sentence epoch to avoid edge artefacts. From this, we derived power estimates from individually defined (i.e., based on participants’ IAF values) theta activity from the start to end of each sentence stimulus, electrode, and from the baseline and delayed testing sessions.

Finally, in order to determine whether changes in neural activity between the sleep and wake conditions were truly oscillatory, we used the irregular-resampling auto-spectral analysis toolbox (IRASA v1.0; (Wen & Liu, 2016) to estimate the 1/ƒ power-law exponent characteristic of background spectral activity, which was used as a covariate in EEG-based statistical models.

#### Sleep parameters and sleep EEG analyses

Sleep data were scored by two sleep technicians (Z.R.C and S.C.) according to standardised criteria (Berry et al., 2012) using Compumedics Profusion 3 software (Melbourne, Australia). The EEG was viewed with a high-pass filter of 0.3 Hz and a low-pass filter of 35 Hz. The following sleep parameters were calculated: total sleep time, sleep onset latency, wake after sleep onset, time (minutes) and percent of time spent in each sleep stage (N1, N2, N3 and R). The EEG data were re-referenced to linked mastoids and filtered from 0.3 – 30 Hz using a digital phase-true FIR band-pass filter. Data were then epoched into 30s bins and subjected to a multivariate covariance-based artifact rejection procedure. This approach estimates a reference covariance matrix for each sleep stage and rejects epochs that deviate too far from this reference, where deviation is established using Riemannian geometry (Barachant et al., 2013; Barthélemy et al., 2019). Slow oscillation-spindle coupling strength was extracted via the danalyzer toolbox implemented in MATLAB based on published algorithms (Denis et al., 2021).

Briefly, sleep spindles were automatically detected at every electrode during NREM sleep based on individual peak spindle frequencies between 12 – 16 Hz. The raw EEG time series was transformed to the frequency domain by estimating the power spectral density (PSD) of the time series using Welch’s method with 5s windows and 50% overlap. Note that the PSD was calculated on a derivative time series to remove the 1/f component and to make the peak spindles more prominent (Demanuele et al., 2007; Sleigh et al., 2001). For each participant at every channel, spindle peak frequencies were automatically detected. Sleep spindles were then automatically detected using a wavelet decomposition, with the Morlet wavelets generated using participants’ peak spindle frequencies. A thresholding algorithm was then applied to every channel to detect spindles in the narrowband data, with a detected spindle needing to exceed a threshold of six times the median amplitude for a minimum of 400ms.

For SOs, continuous NREM EEG data were bandpass filtered between 0.5 and 4 Hz, with all positive-to-negative zero crossings identified based on published alogrithms (Helfrich et al., 2018; Staresina et al., 2015). Potential SOs were flagged if two such positive-to-negative crossings occurred 0.5 – 2s apart. Peak-to-peak amplitudes for all potential SOs were isolated, and oscillations in the top quartile (i.e., with the strongest amplitudes) at each channel were considered SOs (Helfrich et al., 2018; Staresina et al., 2015).

Slow oscillation-spindle coupling was analyzed at each channel during NREM sleep. Specifically, for each identified spindle, we assessed whether it occurred during an identified SO event. These co-occurring events were deemed coupled, and we quantified the percentage of spindle events that were coupled for each channel. For each coupled event, the instantaneous phase of the SO at the time of the peak spindle amplitude was extracted. SO-spindle coupling was further quantified using the mean SO phase and vector length of coupled events for each channel. Finally, the Rayleigh test for circular non-uniformity with alpha set to .01 was used to evaluate phase preference regularity across participants.

### Statistical analysis

Data were imported into *R* version 4.0.2 (R Core Team, 2020) and analysed using (generalised) linear mixed-effects models fit by restricted maximum likelihood (REML) using *lme4* (Bates, 2010). For the behavioural model, we used a logistic mixed-effects regression, modelling response choice (correct, incorrect) as a binary outcome variable. This model also factored in by-item and by-participant differences by specifying them as random effects on the intercept. The behavioural model took the following form:

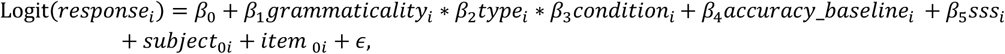

Here, *grammaticality* encodes sentence grammaticality (grammatical, ungrammatical), *type* refers to word order (fixed, flexible), *condition* is sleep versus wake, *baseline* is performance on the baseline (i.e., pre-sleep and -wake) judgement task, and *sss* refers to self-perceived sleepiness estimated from the SSS. Asterisks denote interaction terms, including all subordinate main effects; pluses denote additive terms.

Cluster-based permutation testing (Maris & Oostenveld, 2007) on task-related EEG data was performed in *MATLAB R2022a* (v9.12.0.1884302; The MathWorks, Natick, MA, USA) using the *FieldTrip* toolbox (v20220810; Oostenveld et al., 2011). Baseline-corrected power estimates for each channel and frequency band (theta, alpha, beta) were averaged over the grammaticality factor for both fixed and flexible sentence types. The difference in spectral estimates between fixed and flexible word orders was calculated for each channel and frequency band within-subjects. These difference scores were then contrasted between sleep and wake conditions (thereby testing the interaction between type and condition). Between-subject *t-*statistics were computed using the *ft_statfun_indepsamplesT* function. Channels with *t*-values that exceeded an alpha threshold of .10 were considered as candidates for cluster inclusion. The *t*-values of resolved clusters were then summed and compared to the null distribution of *t*-statistics obtained from 1000 random partitions of the data. The cluster-level statistic was considered significant if it attained a *p*-value < .05.

Following the identification of significant topographical differences in oscillatory power, the following structure was used for the EEG models, where we were interested in predicting behaviour from task-related theta activity, and which did not include trial-based response accuracy:

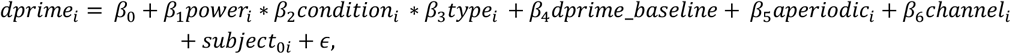

*power* is theta power from the post-sleep and -wake testing session, *condition* is sleep versus wake, and *type* is sentence word order (fixed, flexible). *Baseline* is theta power from the baseline judgement task (pre-sleep and -wake session). a*periodic* refers to the 1/ƒ exponent estimated from the task-related EEG, and *channel* refers to the significant channels isolated from the cluster-based permutation test. Subject was modelled as a random effect on the intercept. d’ was specified as the outcome.

For sleep-related analyses, we first constructed linear mixed-effects model to predict judgement accuracy from the combination of SO-spindle coupling strength, sentence type, sagittality, and laterality, while controlling for baseline (i.e., pre-sleep and -wake) judgement accuracy and sleep stage (N2, N3), with a random intercept of subject. A second linear mixed-effects model was constructed predicting delayed judgement accuracy from anterior task-related theta power, anterior SO-spindle coupling strength and sentence type, while controlling for laterality and baseline judgement accuracy, with random intercepts of subject.

*P*-values for all models were estimated using the *summary* function from the *lmerTest* package, which is based on Satterthwaite’s degrees of freedom (Kuznetsova et al., 2017), while effects were plotted using the package *effects* (Fox & Hong, 2010) and *ggplot2* (Wickham & Wickham, 2016). Post-hoc comparisons for main effects were performed using the *emmeans* package (Lenth et al., 2019). The Holm–Bonferroni method (Holm, 1979) was used to correct for multiple comparisons, while outliers were isolated using Tukey’s method, which identifies outliers as exceeding ± 1.5 × inter-quartile range. Categorical factors were sum-to-zero contrast coded, such that factor level estimates were compared to the grand-mean (Schad et al., 2020). Further, for modelled effects, an 83% confidence interval (CI) threshold was used given that this approach corresponds to the 5% significance level with non-overlapping estimates (Austin & Hux, 2002; MacGregor-Fors & Payton, 2013). In the visualisation of effects, non-overlapping CIs indicate a significant difference at *p* < .05.

## Results

### Sleep supports the consolidation of fixed word order rules

Across testing sessions and grammaticality, participants showed a moderate degree of accuracy for fixed (*M* = 64.00, *SD* = 48.00) and flexible (*M* = 58.00, *SD* = 49.00) word orders, with performance accuracy ranging from 37.18 to 93.75 percent. As shown in Table 1, performance also varied by sentence type, condition, and grammaticality, with the sleep relative to the wake condition performing higher for fixed word orders at delayed testing.

**Table 1.**
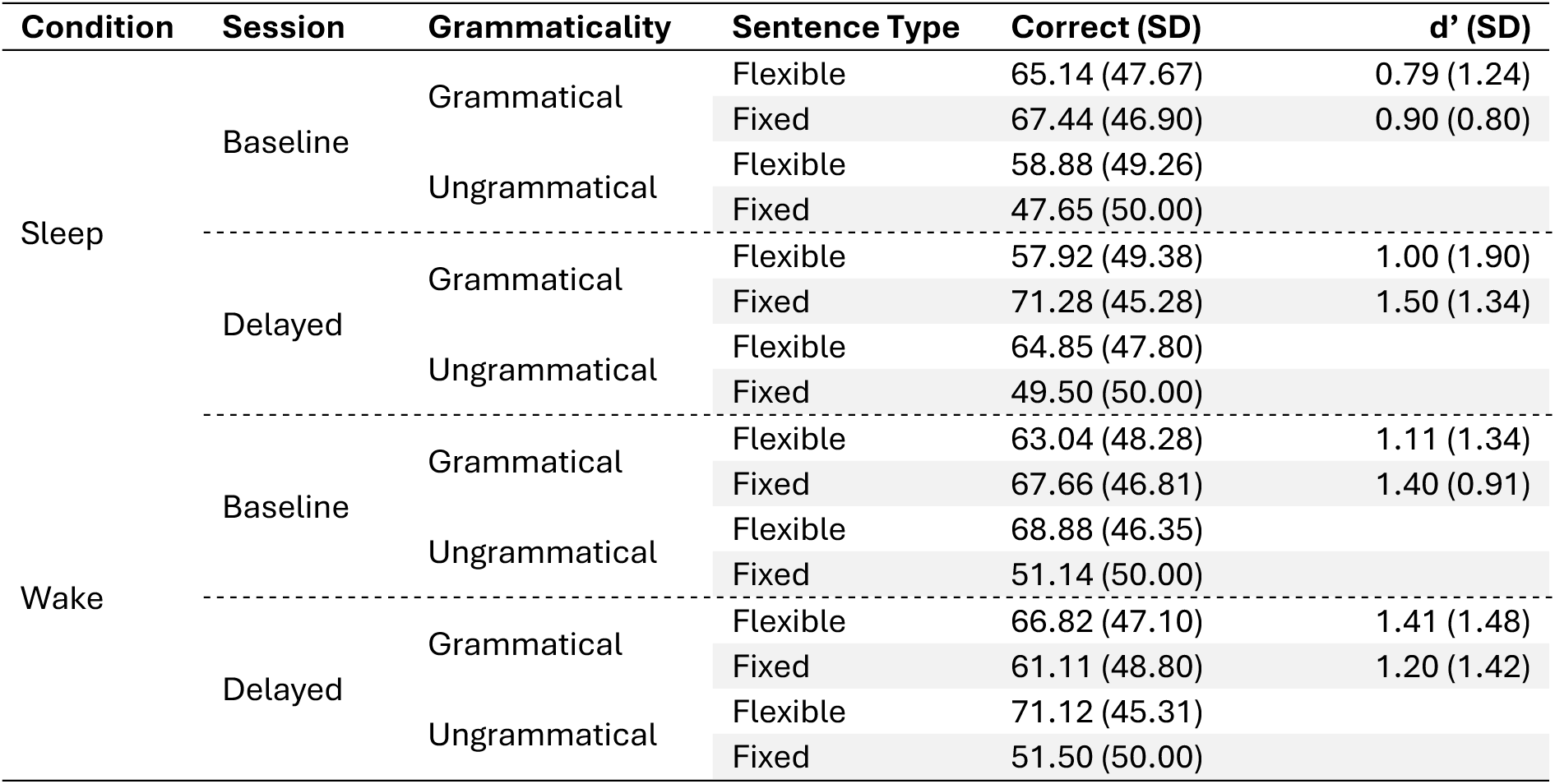
Percent correct and the sensitivity index d’ by condition (sleep, wake), sentence judgement task (baseline, delayed), grammaticality (grammatical, ungrammatical) and sentence type (fixed, flexible). Standard deviations (SD) are given in parentheses.

| Condition | Session | Grammaticality | Sentence Type | Correct (SD) | $d'$ (SD) |
| --- | --- | --- | --- | --- | --- |
| Sleep | Baseline | Grammatical | Flexible | 65.14 (47.67) | 0.79 (1.24) |
|  |  |  | Fixed | 67.44 (46.90) | 0.90 (0.80) |
|  |  | Ungrammatical | Flexible | 58.88 (49.26) |  |
|  |  |  | Fixed | 47.65 (50.00) |  |
|  | Delayed | Grammatical | Flexible | 57.92 (49.38) | 1.00 (1.90) |
|  |  |  | Fixed | 71.28 (45.28) | 1.50 (1.34) |
|  |  | Ungrammatical | Flexible | 64.85 (47.80) |  |
|  |  |  | Fixed | 49.50 (50.00) |  |
| Wake | Baseline | Grammatical | Flexible | 63.04 (48.28) | 1.11 (1.34) |
|  |  |  | Fixed | 67.66 (46.81) | 1.40 (0.91) |
|  |  | Ungrammatical | Flexible | 68.88 (46.35) |  |
|  |  |  | Fixed | 51.14 (50.00) |  |
|  | Delayed | Grammatical | Flexible | 66.82 (47.10) | 1.41 (1.48) |
|  |  |  | Fixed | 61.11 (48.80) | 1.20 (1.42) |
|  |  | Ungrammatical | Flexible | 71.12 (45.31) |  |
|  |  |  | Fixed | 51.50 (50.00) |  |

Generalised linear mixed-effects modelling of single trial response accuracy (controlling for baseline performance) revealed a significant Grammaticality × Sentence Type × Condition interaction (*β* = 0.13, *se* = 0.03, *p* < 0.001; see Figure 4). Holm-Bonferroni adjusted post-hoc comparisons revealed that response accuracy was higher for the sleep relative to wake condition for fixed grammatical (OR = 0.55, *se* = 0.12, *z* = −2.60, *p*_adj_ = 0.03) but not fixed ungrammatical (OR = 0.89, *se* = 0.19, *z* = −0.52, *p*_adj_ = 1.00) word orders.

**Figure 4.**
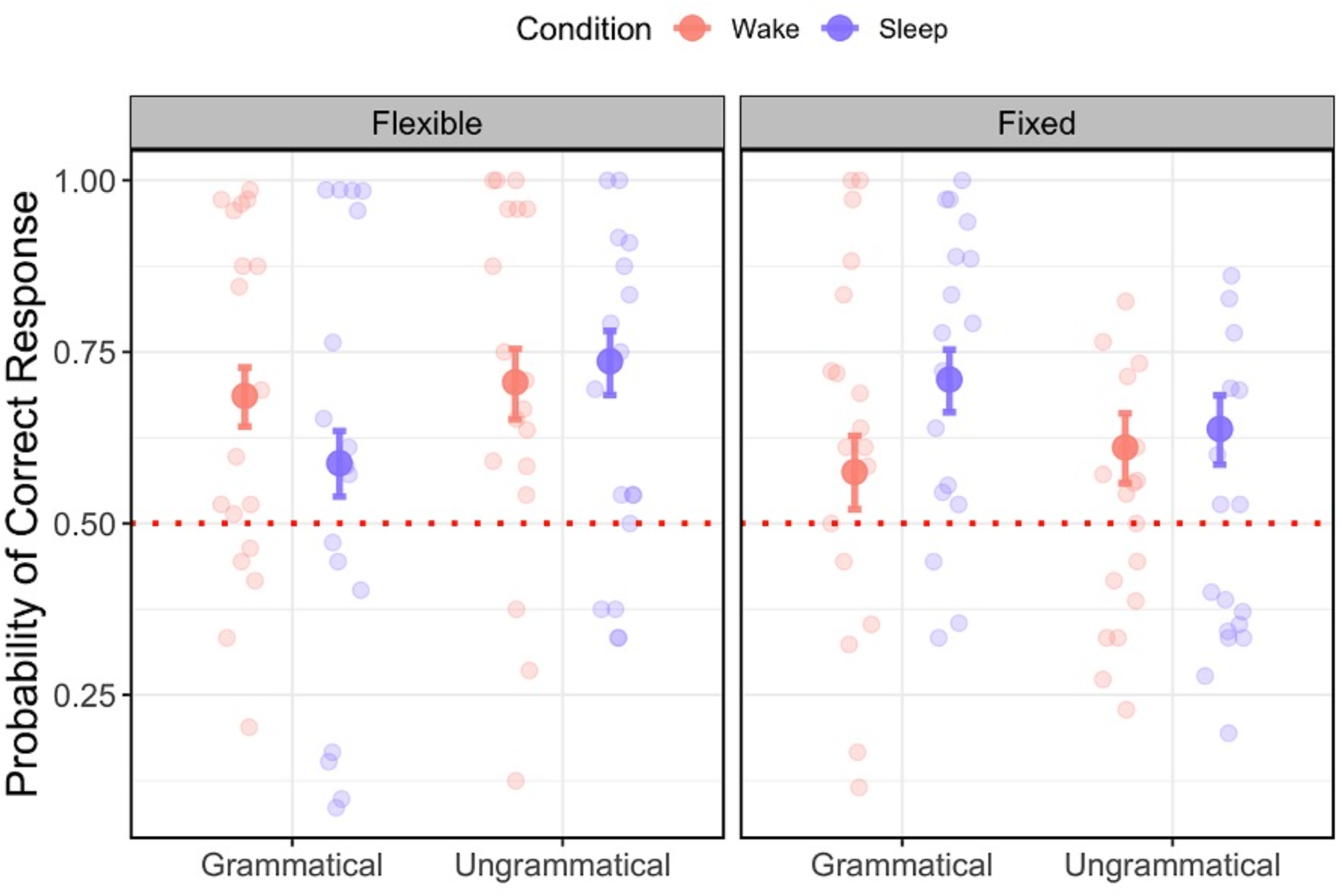
Visualisation of the behavioural results. Relationship between the probability of correct response (y-axis; higher values indicate a higher probability of a correct response), grammaticality (x-axis; grammatical, ungrammatical), sentence type (left column = flexible, right column = fixed), and condition (wake = salmon, sleep = purple). Bars represent the 83% confidence interval around group-level expected marginal mean estimates. Dots represent individual data points per subject for aggregated data.

Response accuracy was also higher in the sleep condition for grammatical fixed relative to grammatical flexible word orders (OR = 0.58, *se* = 0.06, *z* = −4.63, *p*_adj_ < 0.001). The sleep condition also judged flexible over fixed word order sentences as ungrammatical (OR = 1.59, *se* = 0.23, *z* = 3.10, *p*_adj_ = 0.01). These results indicate that sleep may benefit the consolidation of fixed (but not flexible) word order rules, although this pattern may be due to differing response strategies adopted between the sleep and wake conditions. To address this in subsequent analyses, we examine the sensitivity index d’ to account for potential response biases (see Table 1 for d’ values).

### Theta power after sleep is associated with increased memory for fixed, but decreased memory for flexible word order rules

Based on the differences in behavioural performance between the sleep and wake conditions on fixed and flexible word orders, we asked whether task-evoked theta power predicts differences in behaviour across sleep and wake. A non-parametric cluster-based permutation test (see Methods) contrasting Condition (sleep, wake) and Sentence Type (fixed, flexible) revealed a significant difference in baseline-corrected theta power during the delayed session (Monte Carlo *p* = .008; see Figure 5A for topography and demarcation of the cluster). No significant clusters were identified for alpha- or beta-band estimates.

**Figure 5.**
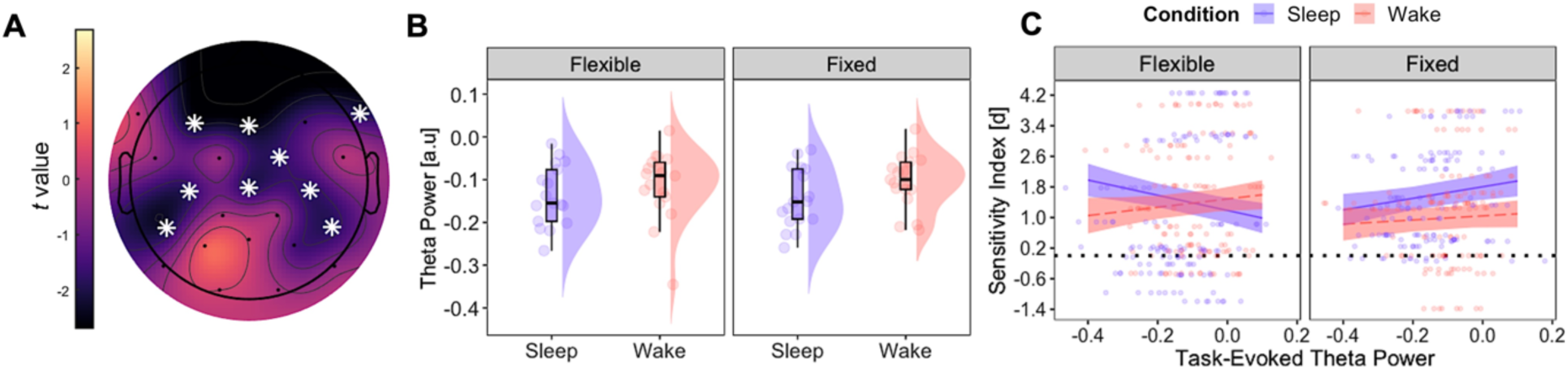
Theta power and judgement accuracy. **(A).** Cluster-based permutation testing on the theta band contrasting differences between Condition (sleep, wake) and Sentence Type (fixed, flexible). Warmer colours denote a higher *t* statistic. Significant channels are indicated by white asterisks. **(B)** Raincloud plots illustrating average theta power over significant channels between sentence type and condition. Positive values on the y-axis denote increased theta power relative to the pre-stimulus interval. **(C)** Modelled effects of task-related theta power (x-axis; higher values indicate increased power) on judgement accuracy (y-axis; higher values indicate better performance) for the sleep and wake conditions (sleep = purple solid line; wake = dashed pink line) for flexible (left facet) and fixed (right facet) sentences. The black dashed line indicates chance-level performance, while the shaded regions indicate the 83% confidence interval. The x-axis reflects theta power estimates, with more negative values reflecting a decrease in power and positive values reflecting an increase in power from the pre-stimulus interval, respectively. Individual data points represent raw (single subject) values.

Given the significant theta-band effects, we constructed a linear mixed-effects model with judgement accuracy (d’) as the outcome and task-related theta power (drawn from the significant cluster identified above), Condition (sleep, wake) and Sentence Type (fixed, flexible) as predictors. This analysis revealed a significant Theta × Condition × Sentence type interaction (*β* = -1.09, *se* = 0.34, *p* = 0.001). Holm-Bonferroni adjusted post-hoc comparisons revealed that for flexible word orders, greater theta synchronisation was associated with poorer judgement accuracy for the sleep but not wake condition. However, the inverse was observed for fixed word order sentences, such that less theta desynchronisation was associated with improved judgement accuracy for the sleep but not wake condition (*β* = -4.70, *se* = 1.10, *p*_adj_ < 0.001). Coupled with the behavioural model, the current analysis demonstrates that sleep preferentially consolidates fixed word order rules at the expense of flexible word order rules, and that this is reflected in task-related theta power. For a visualisation of these effects, see Figure 5C. For time-frequency and power spectral density plots for the sleep and wake conditions across fixed and flexible word orders, see Figures 6 and 7, respectively.

**Figure 6.**
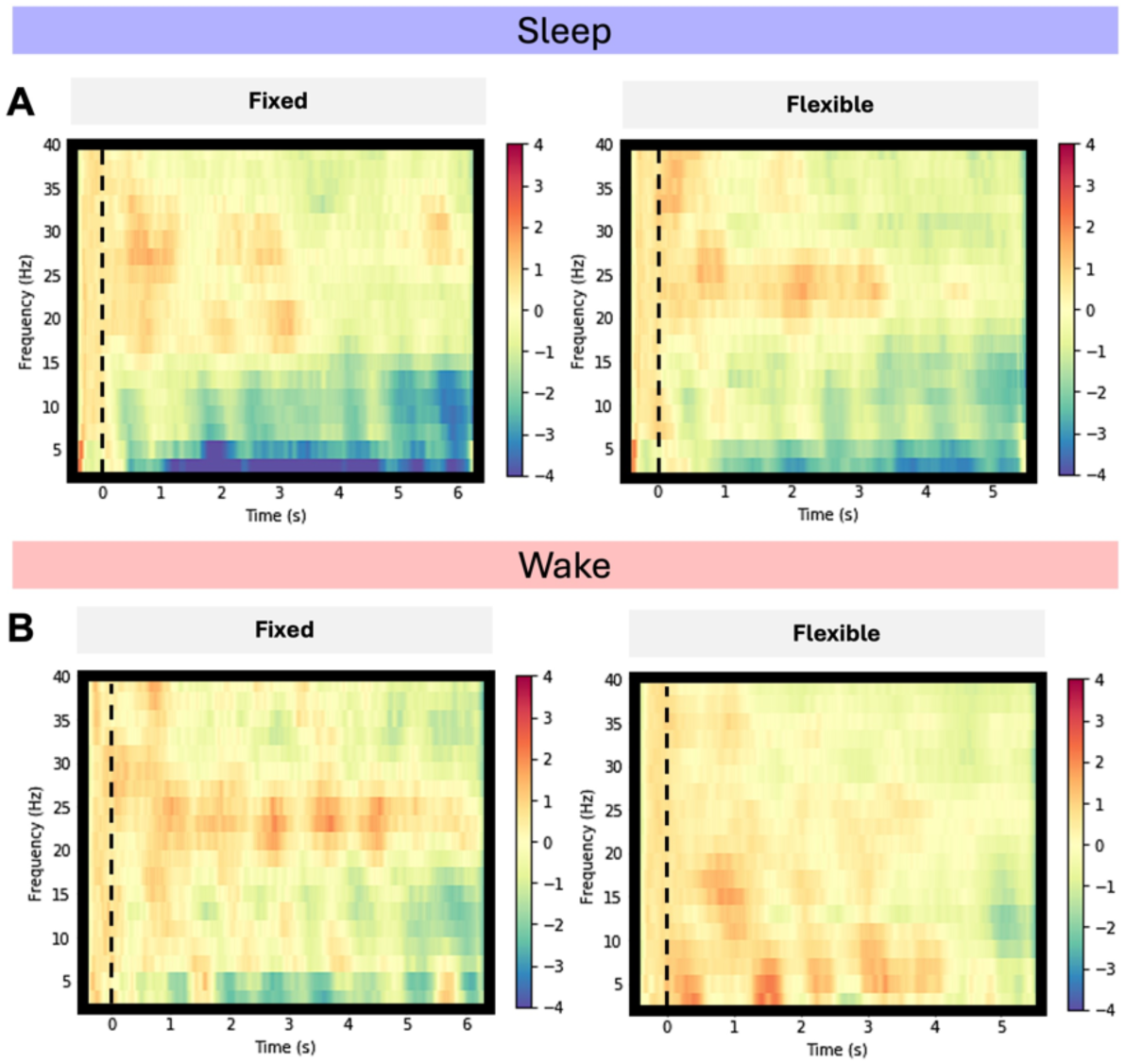
Differences in time-frequency activity between sleep and wake, and fixed and flexible word orders. Time frequency plots for the sleep (top) and wake (bottom) conditions for fixed (left column) and flexible (right column) word order sentences. Time is presented on the x-axis (dashed vertical bar represents sentence onset), while frequency is presented on the y-axis. Warmer colours denote an increase in power relative to the pre-stimulus period, while cooler colours represent a decrease in power. The z-scale is in arbitrary units.

**Figure 7.**
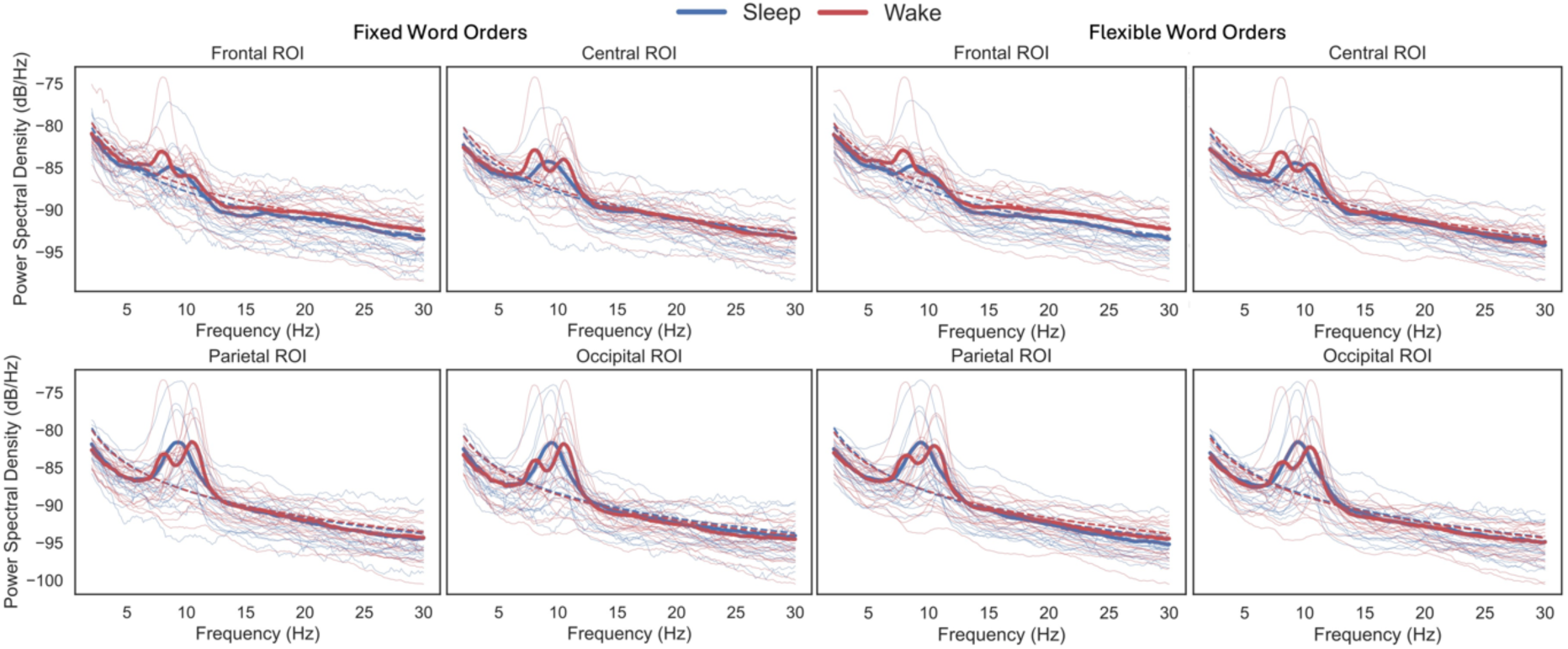
Power spectral density plots for the sleep (blue) and wake (red) conditions for frontal, central, parietal, and occipital regions of interest. Fixed word order sentences are on the left, while flexible word orders are on the right. The solid red and blue lines represent the mean power spectral density for the wake and sleep conditions, respectively, while the dashed lines represent the aperiodic (1/f) power law. Individual lines represent individual participant power spectral densities.

### SO-spindle coupling is predictive of memory for fixed but not flexible word order rules

Having observed differences between the sleep and wake conditions on the relationship between task-related theta activity and behavioural performance, a logical next step was to test whether behavioural performance for fixed word order rules is associated with SO-spindle coupling. Based on previous work (e.g., Helfrich et al., 2018; Mikutta et al., 2019), we focussed on the coupling strength, measured as the mean vector length of spindle phase during coupled SO-spindle events (for a summary of typical sleep parameters and their correlation with d’, see Table 2). There was a significant non-uniform distribution for the precise SO phase during peak spindle activity (*p* ≤ 0.001; Rayleigh test). In predicting behavioural performance, mixed-effects modelling revealed a significant Coupling Strength × Sentence Type × Sagittality interaction (*β* = 3.05, *se* = 0.97, *p* = 0.002). Pairwise contrasts further revealed that this effect was largest anteriorly for fixed sentences (*β* = 6.85, *se* = 2.01, *p*_adj_ < 0.001; Figure 8B), but nonsignificant in central (*β* = -0.75, *se* = 2.62, *p*_adj_ = 0.77) and posterior regions (*β* = -3.90, *se* = 3.47, *p*_adj_ = 0.26). Also note that while stronger SO-spindle coupling predicted improved judgement accuracy for fixed word order sentences, the inverse relationship was present for flexible word order sentences. Figure 8 illustrates an exemplary full-night spectrogram, distribution of SO-spindle coupling strength across channels, as well as exemplar single subject and group level comodulagrams and preferred phase of SO-spindle coupling for NREM sleep. For a summary of sleep microarchitecture characteristics, see Table 3.

**Figure 8.**
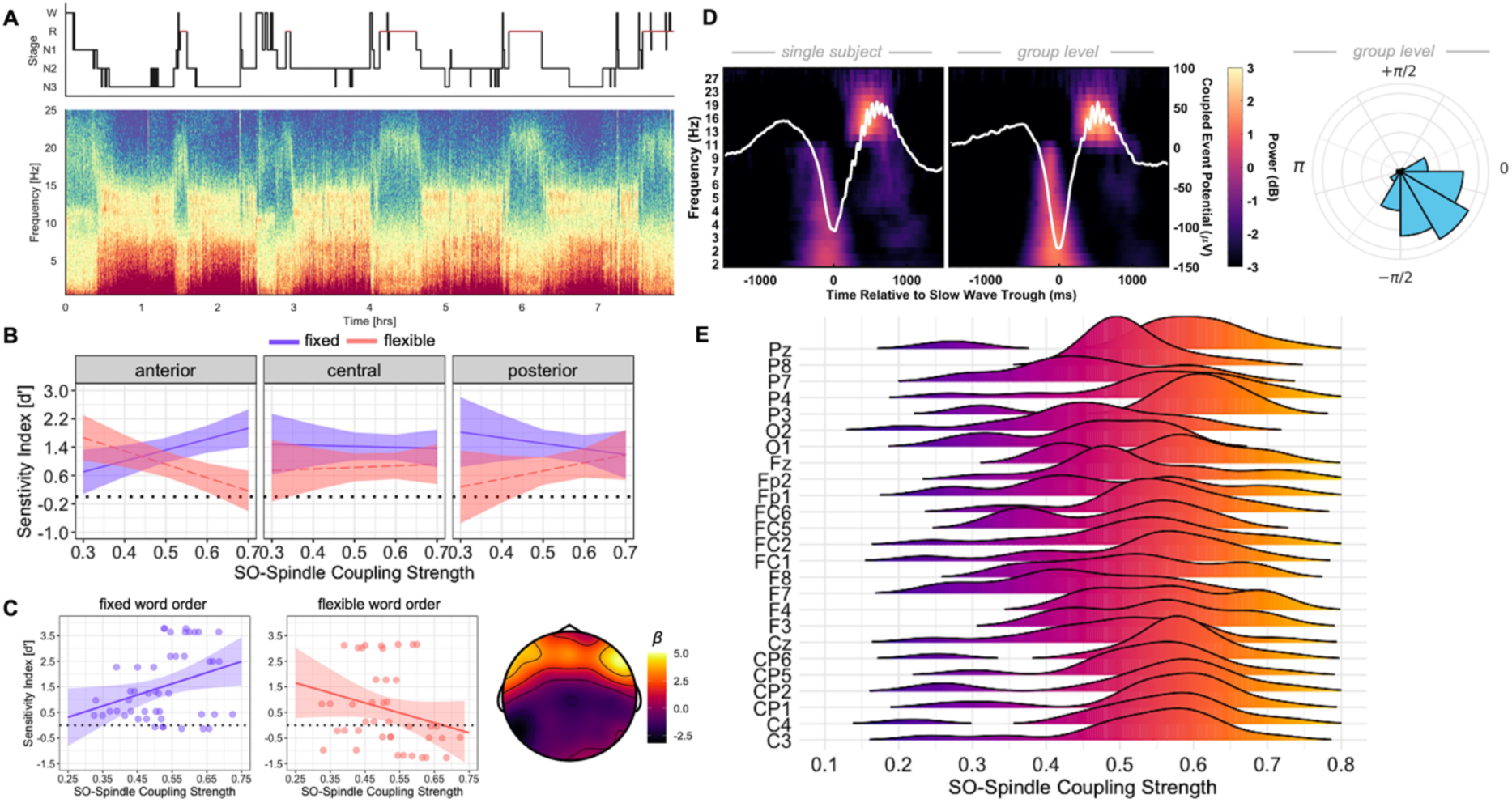
Sleep neurophysiology metrics and relationship between phase amplitude coupling and judgement accuracy. **(A)** Hypnogram and full-night multi-taper spectrogram for a single participant from channel Cz. **(B)** Modelled effects from the linear mixed-effects regression of SO-spindle coupling strength (x-axis; higher values indicate stronger coupling) on judgement accuracy (y-axis; higher values indicate better performance) for fixed and flexible word order sentences (fixed = purple solid line; flexible = dashed pink line) across levels of anterior (left), central (middle) and posterior (right) regions. The black dashed line indicates chance-level performance, while the shaded regions indicate the 83% confidence interval. **(C)** Scatterplot indicating the relationship between judgement accuracy (y-axis; higher values denote better memory performance) and SO-spindle coupling strength (x-axis; higher values denote stronger coupling) for flexible (left) and fixed (right) word order sentences across anterior channels. The topoplot visualises the beta coefficient from the SO-spindle coupling strength × sentence type interaction, with higher values/warmer colours denoting a stronger interaction coefficient. **(D)** Single-subject and group-level average time-frequency response of all SOs coupled to a spindle (−1200 to 1200ms, centred on the trough of the SO), with the time-domain averaged SO overlaid. To the right is the preferred phase of SO-spindle coupling for NREM sleep. Note that 0 represents the peak of the SO. **(E)** Ridge plot illustrating the distribution of SO-spindle coupling strength (x-axis; higher values indicate stronger coupling) across channels (y-axis).

**Table 2.** Descriptive statistics for sleep parameters and correlations with the difference between d’ at delayed and baseline testing for fixed and flexible word order sentences.

| Sleep Parameter | Mean Minutes (SD) | % in Stage (SD) | Correlations with $d'$ (Delayed – Baseline) | | | |
| --- | --- | --- | --- | --- | --- | --- |
|  |  |  | Fixed |  | Flexible |  |
| | | | $r$ | $p$ | $r$ | $p$ |
| TST | 400.00 (67.02) |  | -.44 | .42 | .30 | .96 |
| SOL | 15.23 (12.23) |  | .45 | .42 | -.47 | .35 |
| WASO | 52.64 (55.60) |  | .41 | .42 | -.19 | 1.00 |
| N1 | 38.05 (29.47) | 10.05 (8.21) | .12 | 1.00 | .10 | 1.00 |
| N2 | 196.30 (46.29) | 49.52 (10.36) | .26 | .93 | .33 | .95 |
| SWS | 104.23 (42.27) | 25.84 (9.60) | .02 | 1.00 | -.48 | .35 |
| REM | 61.30 (39.39) | 14.57 (8.56) | -.46 | .42 | .04 | 1.00 |

**Note.** SD = standard deviation. TST = total sleep time; SOL = sleep onset latency; WASO = wake after sleep onset; N1 = stage 1; N2 = stage 2; SWS = slow wave sleep; REM = rapid eye movement sleep. Significance values are Holm-Bonferroni corrected (Holm, 1979).

**Table 3.** NREM slow oscillation-spindle coupling characteristics for frontal, central, and parietal channels.

| Channel | Coupling Strength | Phase | % Coupled | *n* Coupled | *n* Uncoupled | Coupled Density | Uncoupled Density |
| --- | --- | --- | --- | --- | --- | --- | --- |
| C3 | 0.53 (0.11) | -0.27 (0.29) | 16.95 (4.15) | 399.34 (161.21) | 1934.81 (651.20) | 1.33 (0.38) | 6.55 (1.50) |
| C4 | 0.55 (0.11) | -0.24 (0.32) | 17.36 (4.58) | 397.81 (169.80) | 1870.47 (660.39) | 1.32 (0.39) | 6.33 (1.56) |
| F7 | 0.44 (0.11) | 0.08 (0.73) | 18.39 (4.88) | 317.75 (133.49) | 1422.44 (514.52) | 1.09 (0.38) | 4.81 (1.38) |
| F8 | 0.52 (0.11) | -0.14 (0.29) | 18.78 (5.17) | 333.47 (152.94) | 1425.06 (492.26) | 1.13 (0.42) | 4.81 (1.22) |
| P3 | 0.58 (0.11) | -0.34 (0.26) | 17.36 (3.65) | 435.38 (166.33) | 2038.84 (589.56) | 1.45 (0.35) | 6.93 (1.29) |
| P4 | 0.57 (0.12) | -0.39 (0.23) | 17.11 (4.53) | 414.84 (174.65) | 1972.50 (619.66) | 1.37 (0.39) | 6.69 (1.40) |

**Note:** % Coupled = percent of spindles coupled to an SO; *n* Coupled = total number of coupled spindles to SOs; *n* Uncoupled = total number of uncoupled spindles; Coupling Density = average number of coupled spindles to SOs per 30s epoch; Uncoupled Density = average number of uncoupled spindles per 30s epoch. Standard deviations are provided in parentheses.

### Frontal SO-spindle coupling and task-evoked theta power interact to predict judgement accuracy

Having shown that SO-spindle coupling is associated with improved judgement accuracy for fixed word orders, and judgement accuracy is tracked by task-related theta power, we examined whether frontal theta power interacts with frontal SO-spindle coupling strength to predict judgement accuracy. A mixed-effects model regressing SO-spindle coupling strength, task-based theta power, sagittality (anterior, central, posterior), and sentence type (fixed, flexible) onto judgement accuracy revealed a significant three-way interaction between SO-spindle coupling strength, task-based theta power and sentence type (*β* = -41.60, *se* = 16.70, *p* = 0.01). As illustrated in Figure 9, high anterior task-based theta power and stronger anterior SO-spindle coupling was positively associated with delayed judgement accuracy for fixed but not flexible word order sentences. This finding links frontal neural activity in the sleeping and waking brain to predict higher-order language learning.

**Figure 9.**
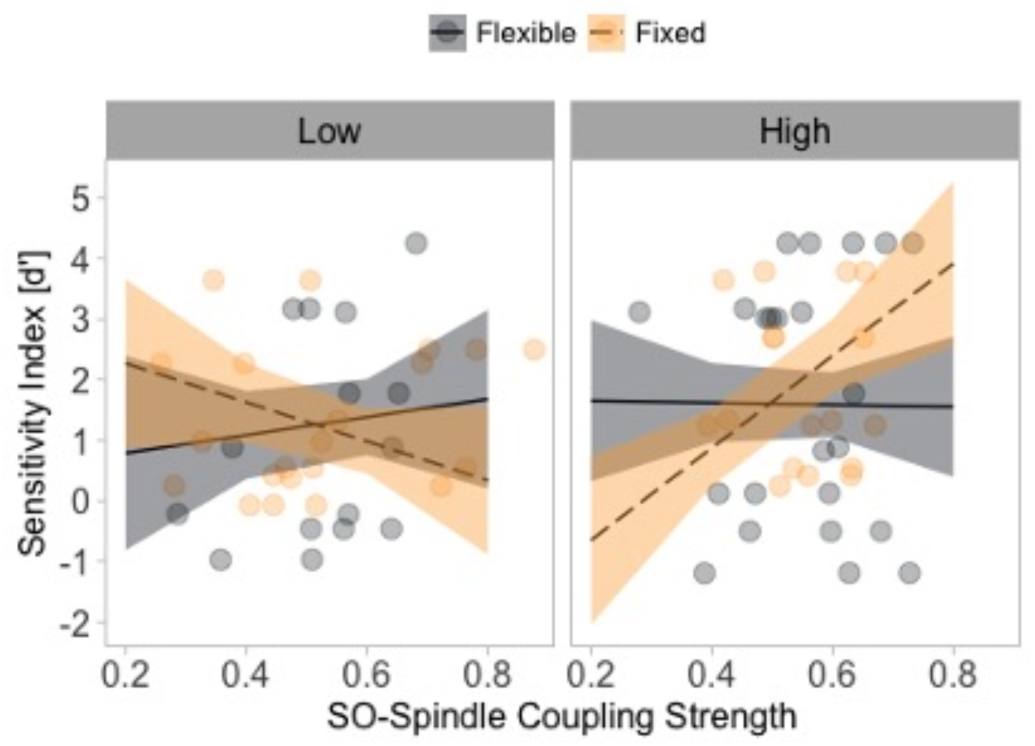
The interaction between task-related theta power and SO-spindle coupling strength predicts judgement accuracy. Delayed judgement accuracy (y-axis; higher values denote higher accuracy), SO-spindle coupling strength (x-axis; higher values denote stronger coupling) and task-related theta power (facetted; low and high contrast for plotting purposes only) averaged across anterior channels. Fixed sentences are colour coded in yellow, while flexible sentences are colour coded in gray.

## Discussion

Coordination between SOs and sleep spindles is hypothesised to provide an optimal temporal receptive window for hippocampal-cortical communication during sleep (Helfrich et al., 2019; Staresina et al., 2015) in the support of memory consolidation. Here, we show that the beneficial effect of SO-spindle coupling on memory extends to sentence-level regularities. Behaviourally, we demonstrated that a period of sleep compared to an equivalent period of wake benefits the consolidation of fixed relative to flexible word order rules, and that this effect is modulated by the strength of coupling between spindles and SOs. Our results further reveal that SO-spindle coupling correlates with changes in task-evoked theta activity during sentence processing. Interestingly, participants in the sleep condition exhibited overall less theta power at delayed testing relative to the wake condition; however, less theta desynchronisation was associated with improved judgement accuracy for fixed word orders in the sleep group. Lastly, we reveal that the interaction between frontal SO-spindle coupling, and task-related frontal theta power predicts improved judgement accuracy for fixed but not flexible word order rules. In sum, our results establish converging behavioural and neurophysiological evidence for a role of NREM SO-spindle coupling and task-related theta activity as signatures of successful memory consolidation and retrieval in the context of higher-order language learning

### Beyond single word learning: a role for sleep in consolidating word order rules

Using a complex modified miniature language paradigm (Cross et al., 2021), we demonstrated that a period of sleep facilitates the extraction of fixed relative to flexible word order rules. Importantly, the key distinction between these word order permutations is that successful interpretation of fixed word order sentences relates to the sequential position of the noun phrases and verb (i.e., the first noun phrase is invariably the actor, and the sentence is verb-final). By contrast, successful interpretation of flexible word order sentences depends more heavily on the animacy of the nouns. As such, fixed word order sentences, requiring a more sequential order-based interpretation and are more compatible with an English word-order-based processing strategy (Bornkessel & Schlesewsky, 2006; Bornkessel-Schlesewsky et al., 2015; MacWhinney et al., 1984). Critically, this sleep-based enhancement for fixed word order rules was predicted by stronger SO-spindle coupling (Figure 8F).

Sleep-related memory effects are proposed to be biased toward stimuli following temporal or sequence-based regularities compared to relational information (for review, see Lerner & Gluck 2019). This is posited to occur via the hippocampal complex encoding temporal occurrences of sensory input (Durrant et al., 2011), which are replayed during SWS, potentially via SO-spindle coupling (e.g., Navarrete et al., 2020; Solano et al., 2020). Here, we provide evidence supporting this account. Specifically, sleep-based consolidation of higher order language may favor sequence-based regularities, with mechanisms of sleep-related memory consolidation generalizing fixed over flexible word order rules, indexed by task-related theta activity.

It is important to note, however, that our sample of participants were native monolingual speakers, and as such, may have preferentially consolidated fixed word order rules at the expense of flexible rules. While behavioral work demonstrates sentence-level preferences of grammatical rules that are analogous to learners’ native languages (e.g., Cross et al., 2021), less is known regarding the neural underpinnings of this phenomenon. We now turn to how the neurobiological processes underpinning the beneficial effect of SO-spindle coupling on memory consolidation extends to higher order language learning.

### Slow oscillation-spindle coupling as a marker of sleep-associated memory consolidation and higher-order language learning

Coupling between SOs and spindles predicts successful overnight memory consolidation (Hahn et al., 2020, 2022; Helfrich et al., 2018; Mikutta et al., 2019). However, these studies often use old-new paradigms with single words (e.g., Helfrich et al., 2018; Mikutta et al., 2019) or word-image pairs (e.g., Muehlroth et al., 2019). Here, we found that the generalisation of sequence-based (or fixed word order) rules is facilitated by the strength of NREM SO-spindle coupling. Mechanistically, during SWS, the cortex is synchronised during the up state of the SO, allowing effective interregional communication, particularly between the prefrontal cortex and hippocampal complex (Helfrich et al., 2019). It is during this SO up-state that spindles induce an influx of Ca^2+^ into excitatory neurons, enabling synaptic plasticity and the generalisation and stabilisation of memory traces (Niethard et al., 2018). Here we revealed that the interaction between these cardinal markers of sleep-related memory processing extend to sentence-level regularities. This finding also accords with previous work examining not only NREM sleep and language learning (Batterink et al., 2014; Mirković & Gaskell, 2016; Schreiner & Rasch, 2017), but also REM (De Koninck et al., 1989, 1990; Thompson et al., 2021). For example, the interaction between time spent in NREM and REM modulates the amplitude of language-related ERPs (N400, late positivity) during the processing of novel grammatical rules (Batterink et al., 2014), while percent of time spent in REM is predictive of French learning in a naturalistic multi-week program (De Koninck et al. 1989, 1990). By demonstrating sleep-related consolidation effects for linguistic stimuli of varying complexity, these findings have begun to establish a link between sleep-related memory consolidation of various aspects of language (Rasch, 2017). Building on this foundational work, we have provided empirical evidence supporting a link between oscillatory-based models of hippocampo-cortical memory consolidation and sentences-level learning, and how this effect manifests in on-task oscillatory theta activity. In the following, we discuss how SO-spindle coupling, as a marker of sleep-associated memory consolidation, modulates task-related oscillatory activity and how these interactions affect sentence processing.

### Task-related theta oscillations index successful memory consolidation of complex linguistic rules

Theta is the dominant frequency in the hippocampal complex and surrounding structures during wake (Covington & Duff, 2016; Duff & Brown-Schmidt, 2012). Oscillations in this frequency range are critical for associative memory formation and coordinating hippocampal-cortical interactions, having been related to associative memory formation (Tort et al., 2009), tracking sequential rules (Crivelli-Decker et al., 2018) and predicting words based on contextual linguistic information (Corcoran et al., 2023; Piai et al., 2016). In the sleep and memory literature, increased theta power has been reported for successfully remembered items, interpreted as reflecting a stronger memory trace induced by sleep-based consolidation. Here, we observed that less theta desynchronisation relative to the pre-stimulus interval predicted higher sensitivity for fixed word order rules after a 12hr delay period, and that the effect of theta on fixed word order processing was more pronounced in the sleep relative to wake condition. This finding accords with the general memory literature, possibly reflecting the binding of linguistic items in a sequence to generate a coherent sentential percept.

We also observed that frontal NREM SO-spindle coupling, and task-related theta power interacted to predict improved delayed judgement accuracy for fixed but not flexible word order rules. In line with systems consolidation theory (Born & Wilhelm, 2012), NREM oscillatory activity contributes to the consolidation of newly encoded memory representations, which may manifest in stronger theta power during retrieval, indicating a stronger neocortical memory trace (Schreiner & Rasch, 2015), reflected in improved sensitivity to fixed word order rules.

### Future directions and concluding remarks

Future studies may include groups in AM-PM (12h Wake), PM-AM (12h Sleep), PM-PM (24h Sleep early) and AM-AM (24h Sleep late), as recommended by Nemeth et al. (2024). We did, however, model participants’ sleepiness levels and the 1/ƒ exponent in our statistical analyses, which partially controlled for potential time-of-day effects. Further, the evidence presented here is correlational and neuroanatomical inferences are unable to be drawn based on scalp-recorded EEG. However, this is the first study to relate sleep-based memory consolidation mechanisms (i.e., SO-spindle coupling) to online sentence-level oscillatory activity, and as such, has set the foundation for future work using techniques with greater spatial-temporal resolution. For example, electrocorticography and stereoelectroencephalography would allow for a better characterization of task-evoked cortical dynamics and SO-spindle coupling between cortical regions and the hippocampal complex, respectively (e.g., Helfrich et al., 2018, 2019). This approach would be complemented by demonstrating a selective reinstatement of memory traces during SO-spindle coupling using representational similarity analysis (Zhang et al., 2018). Identifying stimulus-specific representations during the encoding of sentence-level regularities and tracking the replay of stimulus activity related to SO-spindle coupling events would further demonstrate the critical role of sleep-based oscillatory mechanisms on higher-order language learning. Comparisons between sleep-related consolidation effects on language-specific and non-language but related tasks (i.e., statistical learning tasks) in the same group of participants would also further establish the role of sleep in higher-order language learning.

In addition to representational similarity analyses, we suggest that research examine different baselining approaches to task-related differences in theta activity in conditions of sleep and wake. Here, we adopted a conventional baselining approach of subtracting theta power from the pre-stimulus interval from the stimulus period. In doing so, we observed that the sleep group had greater theta desynchronization than the wake group, but that *less* desynchronization was associated with improved recognition accuracy. From this perspective, it appears that *more* theta power is indeed associated with better memory, but future research should establish whether this effect is driven by a limiting of task-related desynchronization, as we observed, or if a different baselining procedure would reveal an increase in theta power.

Taken together, our results demonstrate that the temporal coupling between NREM SOs and spindles supports the consolidation of complex sentence-level rules. We demonstrated that SO-spindle coupling promotes the consolidation of sequence-based rules and modulates task-evoked theta oscillations previously implicated in language learning (e.g., de Diego-Balaguer et al., 2011; Kepinska et al., 2017) and sentence processing (Vassileiou et al., 2018). Critically, these findings add to models of sleep-based memory consolidation (e.g., Born & Wilhelm, 2012; Lewis & Durrant, 2011) and help characterise how effects of sleep-related oscillatory dynamics on memory manifest in oscillatory activity during complex language-related operations.

## Acknowledgements

We thank Alex Chatburn and Samantha Gray for helpful discussions and feedback on an earlier version of this manuscript. We also thank the research assistants at the Cognitive Neuroscience Laboratory. Particular thanks to Isabella Sharrad, Erica Wilkinson, Nicole Vass, and Angela Osborn for help with data collection. Thank you also to the participants.

## Author Contribution

Conceptualization by Z.C., I.B-S., M.S. and M.J.K. Data curation by Z.C. and L.Z.W. Data pre-processing and analysis by Z.C., S.C., R.F.H., and A.W.C. Writing – original draft by Z.C.; writing – review & editing by all authors; Supervision of project by I.B-S., M.S., M.J.K and R.F.H. Visualization by Z.C.; Funding acquisition by I.B-S. and Z.C.

## References

Austin, Peter C., and Janet E. Hux. 2002. “A Brief Note on Overlapping Confidence Intervals.” Journal of Vascular Surgery 36(1):194–95.

Backus, Alexander R., Jan-Mathijs Schoffelen, Szabolcs Szebényi, Simon Hanslmayr, and Christian F. Doeller. 2016. “Hippocampal-Prefrontal Theta Oscillations Support Memory Integration.” Current Biology 26(4):450–57. doi: 10.1016/j.cub.2015.12.048.

Bakker, Iske, Atsuko Takashima, Janet G. Van Hell, Gabriele Janzen, and James M. McQueen. 2015. “Changes in Theta and Beta Oscillations as Signatures of Novel Word Consolidation.” Journal of Cognitive Neuroscience 27(7):1286–97.

Barachant, Alexandre, Anton Andreev, and Marco Congedo. 2013. “The Riemannian Potato: An Automatic and Adaptive Artifact Detection Method for Online Experiments Using Riemannian Geometry.” Pp. 19–20 in.

Barthélemy, Quentin, Louis Mayaud, David Ojeda, and Marco Congedo. 2019. “The Riemannian Potato Field: A Tool for Online Signal Quality Index of EEG.” IEEE Transactions on Neural Systems and Rehabilitation Engineering 27(2):244–55.

Bastian, Lisa, Anumita Samanta, Demetrius Ribeiro de Paula, Frederik D. Weber, Robby Schoenfeld, Martin Dresler, and Lisa Genzel. 2022. “Spindle–Slow Oscillation Coupling Correlates with Memory Performance and Connectivity Changes in a Hippocampal Network after Sleep.” Human Brain Mapping 43(13):3923–43. doi: 10.1002/hbm.25893.

Bates, Douglas M. 2010. “Lme4: Mixed-Effects Modeling with R.”

Bates, Elizabeth, Antonella Devescovi, and Beverly Wulfeck. 2001. “Psycholinguistics: A Cross-Language Perspective.” Annual Review of Psychology 52(1):369–96.

Batterink, Laura J., and Ken A. Paller. 2017. “Sleep-Based Memory Processing Facilitates Grammatical Generalization: Evidence from Targeted Memory Reactivation.” Sleep and Language Learning 167:83–93. doi: 10.1016/j.bandl.2015.09.003.

Batterink, Laura J., Carmen E. Westerberg, and Ken A. Paller. 2017. “Vocabulary Learning Benefits from REM after Slow-Wave Sleep.” Neurobiology of Learning and Memory 144:102–13. doi: 10.1016/j.nlm.2017.07.001.

Batterink, Laura J., Delphine Oudiette, Paul J. Reber, and Ken A. Paller. 2014. “Sleep Facilitates Learning a New Linguistic Rule.” Neuropsychologia 65:169–79.

Berry, Richard B., Rohit Budhiraja, Daniel J. Gottlieb, David Gozal, Conrad Iber, Vishesh K. Kapur, Carole L. Marcus, Reena Mehra, Sairam Parthasarathy, and Stuart F. Quan. 2012. “Rules for Scoring Respiratory Events in Sleep: Update of the 2007 AASM Manual for the Scoring of Sleep and Associated Events: Deliberations of the Sleep Apnea Definitions Task Force of the American Academy of Sleep Medicine.” Journal of Clinical Sleep Medicine 8(5):597– 619.

Born, Jan, and Ines Wilhelm. 2012. “System Consolidation of Memory during Sleep.” Psychological Research 76:192–203.

Bornkessel-Schlesewsky, Ina, Matthias Schlesewsky, Steven L. Small, and Josef P. Rauschecker. 2015. “Neurobiological Roots of Language in Primate Audition: Common Computational Properties.” Trends in Cognitive Sciences 19(3):142–50.

Bornkessel, Ina, and Matthias Schlesewsky. 2006. “The Extended Argument Dependency Model: A Neurocognitive Approach to Sentence Comprehension across Languages.” Psychological Review 113(4):787.

Brodt, Svenja, Marion Inostroza, Niels Niethard, and Jan Born. 2023. “Sleep—A Brain-State Serving Systems Memory Consolidation.” Neuron 111(7):1050–75. doi: 10.1016/j.neuron.2023.03.005.

Buysse, Daniel J., Charles F. Reynolds III, Timothy H. Monk, Susan R. Berman, and David J. Kupfer. 1989. “The Pittsburgh Sleep Quality Index: A New Instrument for Psychiatric Practice and Research.” Psychiatry Research 28(2):193–213.

Buzsáki, György. 2002. “Theta Oscillations in the Hippocampus.” Neuron 33(3):325–40. doi: 10.1016/S0896-6273(02)00586-X.

Canolty, Ryan T., Erik Edwards, Sarang S. Dalal, Maryam Soltani, Srikantan S. Nagarajan, Heidi E. Kirsch, Mitchel S. Berger, Nicholas M. Barbaro, and Robert T. Knight. 2006. “High Gamma Power Is Phase-Locked to Theta Oscillations in Human Neocortex.” Science 313(5793):1626–28.

Combrisson, Etienne, Marcela Perrone-Bertolotti, Juan LP Soto, Golnoush Alamian, Philippe Kahane, Jean-Philippe Lachaux, Aymeric Guillot, and Karim Jerbi. 2017. “From Intentions to Actions: Neural Oscillations Encode Motor Processes through Phase, Amplitude and Phase-Amplitude Coupling.” NeuroImage 147:473–87.

Corcoran, Andrew W., Phillip M. Alday, Matthias Schlesewsky, and Ina Bornkessel-Schlesewsky. 2018. “Toward a Reliable, Automated Method of Individual Alpha Frequency (IAF) Quantification.” Psychophysiology 55(7):e13064.

Corcoran, Andrew W., Ricardo Perera, Matthieu Koroma, Sid Kouider, Jakob Hohwy, and Thomas Andrillon. 2023. “Expectations Boost the Reconstruction of Auditory Features from Electrophysiological Responses to Noisy Speech.” Cerebral Cortex 33(3):691–708.

Covington, Natalie V., and Melissa C. Duff. 2016. “Expanding the Language Network: Direct Contributions from the Hippocampus.” Trends in Cognitive Sciences 20(12):869–70.

Crivelli-Decker, Jordan, Liang-Tien Hsieh, Alex Clarke, and Charan Ranganath. 2018. “Theta Oscillations Promote Temporal Sequence Learning.” Neurobiology of Learning and Memory 153:92–103.

Cross, Z. R., Corcoran, A. W., Schlesewsky, M., Kohler, M. J., & Bornkessel-Schlesewsky, I. (2022). Oscillatory and aperiodic neural activity jointly predict language learning. Journal of Cognitive Neuroscience, 34(9), 1630–1649.

Cross, Zachariah R., Lena Zou-Williams, Erica M. Wilkinson, Matthias Schlesewsky, and Ina Bornkessel-Schlesewsky. 2021. “Mini Pinyin: A Modified Miniature Language for Studying Language Learning and Incremental Sentence Processing.” Behavior Research Methods 53(3):1218–39. doi: 10.3758/s13428-020-01473-6.

Cross, Zachariah R., Mark J. Kohler, Matthias Schlesewsky, M. G. Gaskell, and Ina Bornkessel- Schlesewsky. 2018. “Sleep-Dependent Memory Consolidation and Incremental Sentence Comprehension: Computational Dependencies during Language Learning as Revealed by Neuronal Oscillations.” Frontiers in Human Neuroscience 12.

Crunelli, Vincenzo, and Stuart W. Hughes. 2010. “The Slow (<1 Hz) Rhythm of Non-REM Sleep: A Dialogue between Three Cardinal Oscillators.” Nature Neuroscience 13(1):9–17. doi: 10.1038/nn.2445.

Davis, Matthew H., and M. Gareth Gaskell. 2009. “A Complementary Systems Account of Word Learning: Neural and Behavioural Evidence.” Philosophical Transactions of the Royal Society B: Biological Sciences 364(1536):3773–3800. doi: 10.1098/rstb.2009.0111.

Demanuele, C., James, C. J., & Sonuga-Barke, E. J. (2007). Distinguishing low frequency oscillations within the 1/f spectral behaviour of electromagnetic brain signals. Behavioral and Brain Functions, 3, 1–14. doi: 10.1186/1744-9081-3-62.

de Diego-Balaguer, Ruth, Lluis Fuentemilla, and Antoni Rodriguez-Fornells. 2011. “Brain Dynamics Sustaining Rapid Rule Extraction from Speech.” Journal of Cognitive Neuroscience 23(10):3105–20.

De Koninck, Joseph, Dominique Lorrain, Günter Christ, Geneviève Proulx, and Daniel Coulombe. 1989. “Intensive Language Learning and Increases in Rapid Eye Movement Sleep: Evidence of a Performance Factor.” International Journal of Psychophysiology 8(1):43–47.

De Koninck, Joseph, G. Christ, G. Hébert, and N. Rinfret. 1990. “Language Learning Efficiency, Dreams and REM Sleep.” Psychiatric Journal of the University of Ottawa: Revue de Psychiatrie de L’universite D’ottawa 15(2):91–92.

Diekelmann, Susanne, Ines Wilhelm, and Jan Born. 2009. “The Whats and Whens of Sleep-Dependent Memory Consolidation.” Sleep Medicine Reviews 13(5):309–21. doi: 10.1016/j.smrv.2008.08.002.

Doppelmayr, M., Klimesch, W., Pachinger, T., & Ripper, B. (1998). Individual differences in brain dynamics: important implications for the calculation of event-related band power. Biological Cybernetics, 79(1), 49–57.

Duff, Melissa C., and Sarah Brown-Schmidt. 2012. “The Hippocampus and the Flexible Use and Processing of Language.” Frontiers in Human Neuroscience 6:69.

Durrant, Simon J., Charlotte Taylor, Scott Cairney, and Penelope A. Lewis. 2011. “Sleep-Dependent Consolidation of Statistical Learning.” Neuropsychologia 49(5):1322–31.

Ferrarelli, F., Huber, R., Peterson, M. J., Massimini, M., Murphy, M., Riedner, B. A., … & Tononi, G. (2007). Reduced sleep spindle activity in schizophrenia patients. American Journal of Psychiatry, 164(3), 483–492.

Fox, John, and Jangman Hong. 2010. “Effect Displays in R for Multinomial and Proportional-Odds Logit Models: Extensions to the Effects Package.” Journal of Statistical Software 32:1–24.

Gramfort, Alexandre, Martin Luessi, Eric Larson, Denis A. Engemann, Daniel Strohmeier, Christian Brodbeck, Roman Goj, Mainak Jas, Teon Brooks, and Lauri Parkkonen. 2013. “MEG and EEG Data Analysis with MNE-Python.” Frontiers in Neuroscience 267.

Hahn, Michael A., Dominik Heib, Manuel Schabus, Kerstin Hoedlmoser, and Randolph F. Helfrich. 2020. “Slow Oscillation-Spindle Coupling Predicts Enhanced Memory Formation from Childhood to Adolescence” edited by S. Haegens and L. L. Colgin. eLife 9:e53730. doi: 10.7554/eLife.53730.

Hahn, Michael A., Kathrin Bothe, Dominik Heib, Manuel Schabus, Randolph F. Helfrich, and Kerstin Hoedlmoser. 2022. “Slow Oscillation–Spindle Coupling Strength Predicts Real-Life Gross-Motor Learning in Adolescents and Adults” edited by S. Haegens and L. L. Colgin. eLife 11:e66761. doi: 10.7554/eLife.66761.

Hautus, Michael J. 1995. “Corrections for Extreme Proportions and Their Biasing Effects on Estimated Values of D′.” Behavior Research Methods, Instruments, & Computers 27:46–51.

Helfrich, Randolph F., Bryce A. Mander, William J. Jagust, Robert T. Knight, and Matthew P. Walker. 2018. “Old Brains Come Uncoupled in Sleep: Slow Wave-Spindle Synchrony, Brain Atrophy, and Forgetting.” Neuron 97(1):221–30.

Helfrich, Randolph F., Janna D. Lendner, Bryce A. Mander, Heriberto Guillen, Michelle Paff, Lilit Mnatsakanyan, Sumeet Vadera, Matthew P. Walker, Jack J. Lin, and Robert T. Knight. 2019. “Bidirectional Prefrontal-Hippocampal Dynamics Organize Information Transfer during Sleep in Humans.” Nature Communications 10(1):3572. doi: 10.1038/s41467-019-11444-x.

Holm, Sture. 1979. “A Simple Sequentially Rejective Multiple Test Procedure.” Scandinavian Journal of Statistics 65–70.

Hyvarinen, Aapo. 1999. “Fast and Robust Fixed-Point Algorithms for Independent Component Analysis.” IEEE Transactions on Neural Networks 10(3):626–34.

Isbilen, Erin S., Stewart M. McCauley, and Morten H. Christiansen. 2022. “Individual Differences in Artificial and Natural Language Statistical Learning.” Cognition 225:105123. doi: 10.1016/j.cognition.2022.105123.

Jacobs, Joshua, Grace Hwang, Tim Curran, and Michael J. Kahana. 2006. “EEG Oscillations and Recognition Memory: Theta Correlates of Memory Retrieval and Decision Making.” NeuroImage 32(2):978–87. doi: 10.1016/j.neuroimage.2006.02.018.

James, Emma, M. Gareth Gaskell, Anna Weighall, and Lisa Henderson. 2017. “Consolidation of Vocabulary during Sleep: The Rich Get Richer?” Neuroscience & Biobehavioral Reviews 77:1–13.

Kavanau, JL. 1997. “Memory, Sleep and the Evolution of Mechanisms of Synaptic Efficacy Maintenance.” Neuroscience 79(1):7–44.

Kepinska, Olga, Ernesto Pereda, Johanneke Caspers, and Niels O. Schiller. 2017. “Neural Oscillatory Mechanisms during Novel Grammar Learning Underlying Language Analytical Abilities.” Brain and Language 175:99–110.

Klimesch, Wolfgang. 2012. “Alpha-Band Oscillations, Attention, and Controlled Access to Stored Information.” Trends in Cognitive Sciences 16(12):606–17.

Klinzing, Jens G., Niels Niethard, and Jan Born. 2019. “Mechanisms of Systems Memory Consolidation during Sleep.” Nature Neuroscience 22(10):1598–1610.

Köster, Moritz, Holger Finger, Maren-Jo Kater, Christoph Schenk, and Thomas Gruber. 2017. “Neuronal Oscillations Indicate Sleep-Dependent Changes in the Cortical Memory Trace.” Journal of Cognitive Neuroscience 29(4):698–707.

Kuznetsova, Alexandra, Per B. Brockhoff, and Rune HB Christensen. 2017. “lmerTest Package: Tests in Linear Mixed Effects Models.” Journal of Statistical Software 82:1–26

Lenth, Russell, H. Singmann, J. Love, P. Buerkner, and M. Herve. 2019. “Package ‘Emmeans.’”

Lerner, Itamar, and Mark A. Gluck. 2019. “Sleep and the Extraction of Hidden Regularities: A Systematic Review and the Importance of Temporal Rules.” Sleep Medicine Reviews 47:39– 50.

Lewis, Penelope A., and Simon J. Durrant. 2011. “Overlapping Memory Replay during Sleep Builds Cognitive Schemata.” Trends in Cognitive Sciences 15(8):343–51.

MacGregor-Fors, Ian, and Mark E. Payton. 2013. “Contrasting Diversity Values: Statistical Inferences Based on Overlapping Confidence Intervals.” PLoS One 8(2):e56794.

MacWhinney, Brian, Elizabeth Bates, and Reinhold Kliegl. 1984. “Cue Validity and Sentence Interpretation in English, German, and Italian.” Journal of Verbal Learning and Verbal Behavior 23(2):127–50.

Maingret, Nicolas, Gabrielle Girardeau, Ralitsa Todorova, Marie Goutierre, and Michaël Zugaro. 2016. “Hippocampo-Cortical Coupling Mediates Memory Consolidation during Sleep.” Nature Neuroscience 19(7):959–64.

Malerba, P., Whitehurst, L. N., Simons, S. B., & Mednick, S. C. (2019). Spatio-temporal structure of sleep slow oscillations on the electrode manifold and its relation to spindles. Sleep, 42(1), zsy197.

Mathôt, Sebastiaan, Daniel Schreij, and Jan Theeuwes. 2012. “OpenSesame: An Open-Source, Graphical Experiment Builder for the Social Sciences.” Behavior Research Methods 44:314–24.

Mikutta, C., Feige, B., Maier, J. G., Hertenstein, E., Holz, J., Riemann, D., & Nissen, C. (2019). Phase-amplitude coupling of sleep slow oscillatory and spindle activity correlates with overnight memory consolidation. Journal of sleep research, 28(6), e12835.

Mirković, Jelena, and M. Gareth Gaskell. 2016. “Does Sleep Improve Your Grammar? Preferential Consolidation of Arbitrary Components of New Linguistic Knowledge.” PloS One 11(4):e0152489.

Muehlroth, Beate E., Myriam C. Sander, Yana Fandakova, Thomas H. Grandy, Björn Rasch, Yee Lee Shing, and Markus Werkle-Bergner. 2019. “Precise Slow Oscillation–Spindle Coupling Promotes Memory Consolidation in Younger and Older Adults.” Scientific Reports 9(1):1940. doi: 10.1038/s41598-018-36557-z.

Navarrete, Miguel, Jules Schneider, Hong-Viet V. Ngo, Mario Valderrama, Alexander J. Casson, and Penelope A. Lewis. 2020. “Examining the Optimal Timing for Closed-Loop Auditory Stimulation of Slow-Wave Sleep in Young and Older Adults.” Sleep 43(6):zsz315.

Németh, D., Gerbier, E., Born, J., Rickard, T., Diekelmann, S., Fogel, S., … & Janacsek, K. (2024). Optimizing the methodology of human sleep and memory research. Nature Reviews Psychology, 3(2), 123–137.

Nicholls, Michael ER, Nicole A. Thomas, Tobias Loetscher, and Gina M. Grimshaw. 2013. “The Flinders Handedness Survey (FLANDERS): A Brief Measure of Skilled Hand Preference.” Cortex 49(10):2914–26.

Nicolas, J., King, B. R., Levesque, D., Lazzouni, L., Coffey, E., Swinnen, S., … & Albouy, G. (2022). Sigma oscillations protect or reinstate motor memory depending on their temporal coordination with slow waves. Elife, 11, e73930.

Niethard, Niels, Hong-Viet V. Ngo, Ingrid Ehrlich, and Jan Born. 2018. “Cortical Circuit Activity Underlying Sleep Slow Oscillations and Spindles.” Proceedings of the National Academy of Sciences 115(39):E9220–29. doi: 10.1073/pnas.1805517115.

Nieuwenhuis, Ingrid LC, Vasiliki Folia, Christian Forkstam, Ole Jensen, and Karl Magnus Petersson. 2013. “Sleep Promotes the Extraction of Grammatical Rules.” PloS One 8(6):e65046.

Oostenveld, Robert, Pascal Fries, Eric Maris, and Jan-Mathijs Schoffelen. 2011. “FieldTrip: Open Source Software for Advanced Analysis of MEG, EEG, and Invasive Electrophysiological Data.” Computational Intelligence and Neuroscience 2011:1–9.

Özkurt, Tolga Esat, and Alfons Schnitzler. 2011. “A Critical Note on the Definition of Phase– Amplitude Cross-Frequency Coupling.” Journal of Neuroscience Methods 201(2):438–43.

Özkurt, Tolga Esat. 2012. “Statistically Reliable and Fast Direct Estimation of Phase-Amplitude Cross-Frequency Coupling.” IEEE Transactions on Biomedical Engineering 59(7):1943–50.

Piai, Vitória, Kristopher L. Anderson, Jack J. Lin, Callum Dewar, Josef Parvizi, Nina F. Dronkers, and Robert T. Knight. 2016. “Direct Brain Recordings Reveal Hippocampal Rhythm Underpinnings of Language Processing.” Proceedings of the National Academy of Sciences 113(40):11366–71. doi: 10.1073/pnas.1603312113.

Purcell, S. M., Manoach, D. S., Demanuele, C., Cade, B. E., Mariani, S., Cox, R., … & Stickgold, R. (2017). Characterizing sleep spindles in 11,630 individuals from the National Sleep Research Resource. Nature communications, 8(1), 15930.

Rasch, Björn, and Jan Born. 2013. “About Sleep’s Role in Memory.” Physiological Reviews 93(2):681–766. doi: 10.1152/physrev.00032.2012.

Rasch, Björn. 2017. “Sleep and Language Learning.” Sleep and Language Learning 167:1–2. doi: 10.1016/j.bandl.2017.02.002.

Romberg, Alexa R., and Jenny R. Saffran. 2010. “Statistical Learning and Language Acquisition.” WIREs Cognitive Science 1(6):906–14. doi: 10.1002/wcs.78.

Santolin, Chiara, and Jenny R. Saffran. 2018. “Constraints on Statistical Learning Across Species.” Trends in Cognitive Sciences 22(1):52–63. doi: 10.1016/j.tics.2017.10.003.

Sattari, N., Whitehurst, L. N., Ahmadi, M., & Mednick, S. C. (2019). Does working memory improvement benefit from sleep in older adults?. Neurobiology of sleep and circadian rhythms, 6, 53–61.

Sauppe, S., Choudhary, K. K., Giroud, N., Blasi, D. E., Norcliffe, E., Bhattamishra, S., … & Bickel, B. (2021). Neural signatures of syntactic variation in speech planning. PLoS Biology, 19(1), e3001038. doi: 10.1371/journal.pbio.3001038

Schad, Daniel J., Shravan Vasishth, Sven Hohenstein, and Reinhold Kliegl. 2020. “How to Capitalize on a Priori Contrasts in Linear (Mixed) Models: A Tutorial.” Journal of Memory and Language 110:104038.

Schreiner, Thomas, and Björn Rasch. 2015. “Boosting Vocabulary Learning by Verbal Cueing during Sleep.” Cerebral Cortex 25(11):4169–79.

Schreiner, Thomas, and Björn Rasch. 2017. “The Beneficial Role of Memory Reactivation for Language Learning during Sleep: A Review.” Sleep and Language Learning 167:94–105. doi: 10.1016/j.bandl.2016.02.005.

Sleigh, J. W., Steyn-Ross, D. A., Steyn-Ross, M. L., Williams, M. L., & Smith, P. (2001). Comparison of changes in electroencephalographic measures during induction of general anaesthesia: influence of the gamma frequency band and electromyogram signal. British journal of anaesthesia, 86(1), 50–58. doi: 10.1093/bja/86.1.50.

Solano, Agustín, Luis A. Riquelme, Daniel Perez-Chada, and Valeria Della-Maggiore. 2020. “Local Coupling between Sleep Spindles and Slow Oscillations Supports the Stabilization of Motor Memories.” BioRxiv 2020–08.

Stanislaw, Harold, and Natasha Todorov. 1999. “Calculation of Signal Detection Theory Measures.” Behavior Research Methods, Instruments, & Computers 31(1):137–49.

Staresina, Bernhard P., Til Ole Bergmann, Mathilde Bonnefond, Roemer van der Meij, Ole Jensen, Lorena Deuker, Christian E. Elger, Nikolai Axmacher, and Juergen Fell. 2015. “Hierarchical Nesting of Slow Oscillations, Spindles and Ripples in the Human Hippocampus during Sleep.” Nature Neuroscience 18(11):1679–86. doi: 10.1038/nn.4119.

Thompson, Kristen, Aaron Gibbings, James Shaw, Laura Ray, Gilles Hébert, Joseph De Koninck, and Stuart Fogel. 2021. “Sleep and Second-Language Acquisition Revisited: The Role of Sleep Spindles and Rapid Eye Movements.” Nature and Science of Sleep 1887–1902.

Tort, Adriano BL, Robert W. Komorowski, Joseph R. Manns, Nancy J. Kopell, and Howard Eichenbaum. 2009. “Theta–Gamma Coupling Increases during the Learning of Item– Context Associations.” Proceedings of the National Academy of Sciences 106(49):20942– 47.

Vallat, R., Shah, V. D., & Walker, M. P. (2023). Coordinated human sleeping brainwaves map peripheral body glucose homeostasis. Cell Reports Medicine, 4(7).

Vallat, Raphael, and Matthew P. Walker. 2021. “An Open-Source, High-Performance Tool for Automated Sleep Staging.” Elife 10:e70092.

Vallat, Raphael. 2018. “Pingouin: Statistics in Python.” J. Open Source Softw. 3(31):1026.

Vassileiou, Benedict, Lars Meyer, Caroline Beese, and Angela D. Friederici. 2018. “Alignment of Alpha-Band Desynchronization with Syntactic Structure Predicts Successful Sentence Comprehension.” NeuroImage 175:286–96.

Vyazovskiy, Vladyslav V., and Kenneth D. Harris. 2013. “Sleep and the Single Neuron: The Role of Global Slow Oscillations in Individual Cell Rest.” Nature Reviews Neuroscience 14(6):443–51. doi: 10.1038/nrn3494.

Wen, Haiguang, and Zhongming Liu. 2016. “Separating Fractal and Oscillatory Components in the Power Spectrum of Neurophysiological Signal.” Brain Topography 29:13–26.

Wickham, Hadley, and Hadley Wickham. 2016. “Data Analysis.” Ggplot2: Elegant Graphics for Data Analysis 189–201.

Xie, Xin, F. Sayako Earle, and Emily B. Myers. 2018. “Sleep Facilitates Generalisation of Accent Adaptation to a New Talker.” Language, Cognition and Neuroscience 33(2):196–210. doi: 10.1080/23273798.2017.1369551.

Zhang, Hui, Juergen Fell, and Nikolai Axmacher. 2018. “Electrophysiological Mechanisms of Human Memory Consolidation.” Nature Communications 9(1):4103.

